# Multiscale imaging of cytokinesis

**DOI:** 10.1101/2025.09.24.678330

**Authors:** N. Hümpfer, Z. Marin, A. Zehtabian, K. Bruckmann, D. Spierling, J. Schmoranzer, J. Ries, H. Ewers

## Abstract

After separation of duplicated genetic material, animal cells divide by cytokinesis. This process involves several simultaneous and successive steps of molecular reorganization at cell membrane, cortex and the spindle apparatus as the cleavage furrow ingresses and the spindle is transformed into the intercellular bridge. Dozens of proteins play a role in cytokinesis. However, a comprehensive view of their successive nanoscale reorganizations over cytokinesis remains untractable with current microscopy methods. Here, we developed a multimodal imaging and analysis framework to investigate the relative organization of key cytokinetic molecules over time. We extract cytokinetic hallmarks from live-cell lattice light-sheet and expansion microscopy (ExM) to computationally define pseudotime and align nanoscale ExM images of cytokinesis. We discover a cytokinetic substep and create nanoscale maps of the intercellular bridge and the midbody in 6 stages for 20 involved molecules. We thus provide a framework for nanoscale analysis of cytokinesis, a process fundamental to life.

## Introduction

Cell division is the fundamental hallmark of life. Eukaryotic cells divide by a process called cytokinesis after the duplication of the genetic material. In ophistokonts, cytokinesis (mostly^1^) happens via quick ingression of a cleavage furrow at the site of the spindle, where actomyosin constricts the cell until the former spindle is compressed into an elongated bundle between the two daughter cells. Bundled spindle microtubules form a thinning intercellular bridge (ICB) with a bulbous midbody at the previous spindle midzone. This bundle is then cut close to the midzone followed by final separation of the daughter cell cytoplasms by plasma membrane fission^2^. To execute this complex multistep process, the function of hundreds of molecules is required. While the molecular players of most essential substeps have been identified, the exact sequence and nanoscale orchestration of all required functions remains elusive. This is because the extremely dense and complex, dynamic and amorphous nanoscale midbody structure^3^ and the long timeframe of the abscission process have so far prevented a detailed spatiotemporal investigation of the entire process. Live cell microscopy does not allow for fast and long-term super resolution imaging, even less for multiple components. Multiplexed imaging of hundreds of substeps in still images offers a solution, but requires the generation of pseudotime from the known trajectory of reliable features as in clathrin-coated pits^4^ or the centriole^5^. We here reasoned that the super-resolved temporal progression of cytokinesis can be imaged in similar way for a number of targets by sampling the process in hundreds of multicolor expansion microscopy (ExM)^6^ images, where two markers allow for extraction of the exact timepoint in cytokinetic progression.

By now, more than 150 individual proteins have been identified to play a role in cytokinesis^7–9^. They control a number of functions that are executed sequentially and concomitantly with one another to prime the spindle plane membrane for recruitment of the contractile machinery^10,11^, to form the machinery itself^12–14^ and to connect it to the plasma membrane for cleavage furrow ingression^15–18^. They endocytose the excess membrane that results from reduction of membrane area at the cleavage furrow^19,20^. Others regulate tension of the ICB plasma membrane^21^, check if no chromosomal DNA is left between the dividing cells^22,23^ constrict the spindle microtubules into a tight bundle^24,25^, and assemble the midbody^3^. Dedicated machineries are recruited close to the midbody to sever the microtubule bundle^26–29^, cut midbody actin^30,31^ and finally mediate membrane fission to separate daughter cell cytoplasms via ESCRT (Endosomal Sorting Complex Required for Transport)^28,29,32–34^.

Strikingly, a conserved family of GTPases called septins, discovered as cell-division cycle (cdc) mutants in yeast^35^ and with essential function in mammalian cytokinesis^36–38^, play a role in many of these individual functions. They become recruited to the spindle midzone membrane via PI(4,5)P_2_ (Phosphatidylinositol 4,5-bisphosphate)^39^, interact with non-muscle myosin II^40^ and Anillin at the membrane^41^. They locate to the cleavage furrow^37,42^ in dependence of PIP-kinase1γ^43^. They become phosphorylated by checkpoint kinase AuroraB^44^, locate to the midbody and, in the late stages of cytokinesis, play a role in microtubule severing^36^ and finally ESCRT recruitment^45^. The multitude of roles of septins in cytokinesis is further supported by the observation that Septin2 and Septin9 knockdowns display different cytokinetic defects^46^. We thus reasoned that septin localization might be a good marker for cytokinetic progression.

Here, we thus combined fast, long-term live cell imaging of cytokinesis in lattice light-sheet (LLS) microscopy with computational image analysis of ExM to develop a pseudotime nanoscale spatiotemporal map of cytokinesis. We deduce the temporal development of intracellular bridge and midbody progression from microtubule and septin staining and assemble a multicomponent three-dimensional superresolved map from more than 15 proteins involved in cytokinesis. We identify a new substep in cytokinetic progression and reveal nanoscale reorganization of cortical actomyosin, Septin2, Anillin, the midbody and ESCRT, respectively over time. Our assay provides a much-needed experimental paradigm for the investigation of cytokinesis and other large-scale cellular processes at the nanoscale over time.

## Results

### Superresolution assay of cytokinesis

We here aimed to analyze cytokinesis based on lattice-light sheet live-cell imaging of many cell divisions. We then combined nanoscopic Expansion microscopy^6,47^ imaging with pseudotime assignment via principle component analysis of images from defined stages. For live cell imaging, we used a diploid rat NRK49F cell-line that had eGFP inserted into the Septin2 gene on both chromosomes via TALEN (Transcription Activator-Like Effector Nuclease) genome engineering and was carefully characterized before^48^. We synchronized these cells, added SiR-tubulin^49^ and imaged full cell volumes after release on a lattice light-sheet microscope setup at a framerate of 1 per 5 min (Figure 1A). We observed many complete cell division processes with a negligible amount of failures and could clearly distinguish individual steps in the process according to the Septin2-GFP^EN/EN^ and SiR-tubulin fluorescence (Figures 1B, C and Supplementary Movie 1). According to these time points, we could align the temporal progression of cytokinesis in a total of 129 cells (Figure 1D) and found that constriction had happened about 9.6 ± 2.8 min after the cleavage furrow started to ingress (“round and constricted”, RC) and after 15 ± 3.3 min, cells had spread out again on the substrate (“cell spreading” CS). After 23.3 ± 5.9 min, cells exhibited split septin rings at the ICB (septin “ring split”, RS). Abscission as defined by severing of the microtubule bundle was observed earliest after about half an hour and could be as late as after 3h, with 75% of cells requiring between 30 and 90 min to progress to abscission (Figure 1E-F). We aimed to use this dataset to ask if image features could be used as proxies for the temporal progression of cytokinesis. To do so, we used the thickness of the microtubule bundle in the ICB as an accessible feature. When we extracted microtubule thickness from all cells at all time points, we found that microtubule thickness decreased on average over time (Figure 1G), however not equally in all cells. We concluded that this simple image feature alone could thus not allow to determine the relative position of a single cell image in the temporal progression of cytokinesis. Nevertheless, we reasoned that taken together with the feature-rich images of Septin2-GFP^EN/EN^ fluorescence, the thinning of the stained microtubule bundle could allow the creation of a pseudotime representation of features during cytokinesis using principle component analysis (PCA) if sufficient detail was present.

**Figure 1:**
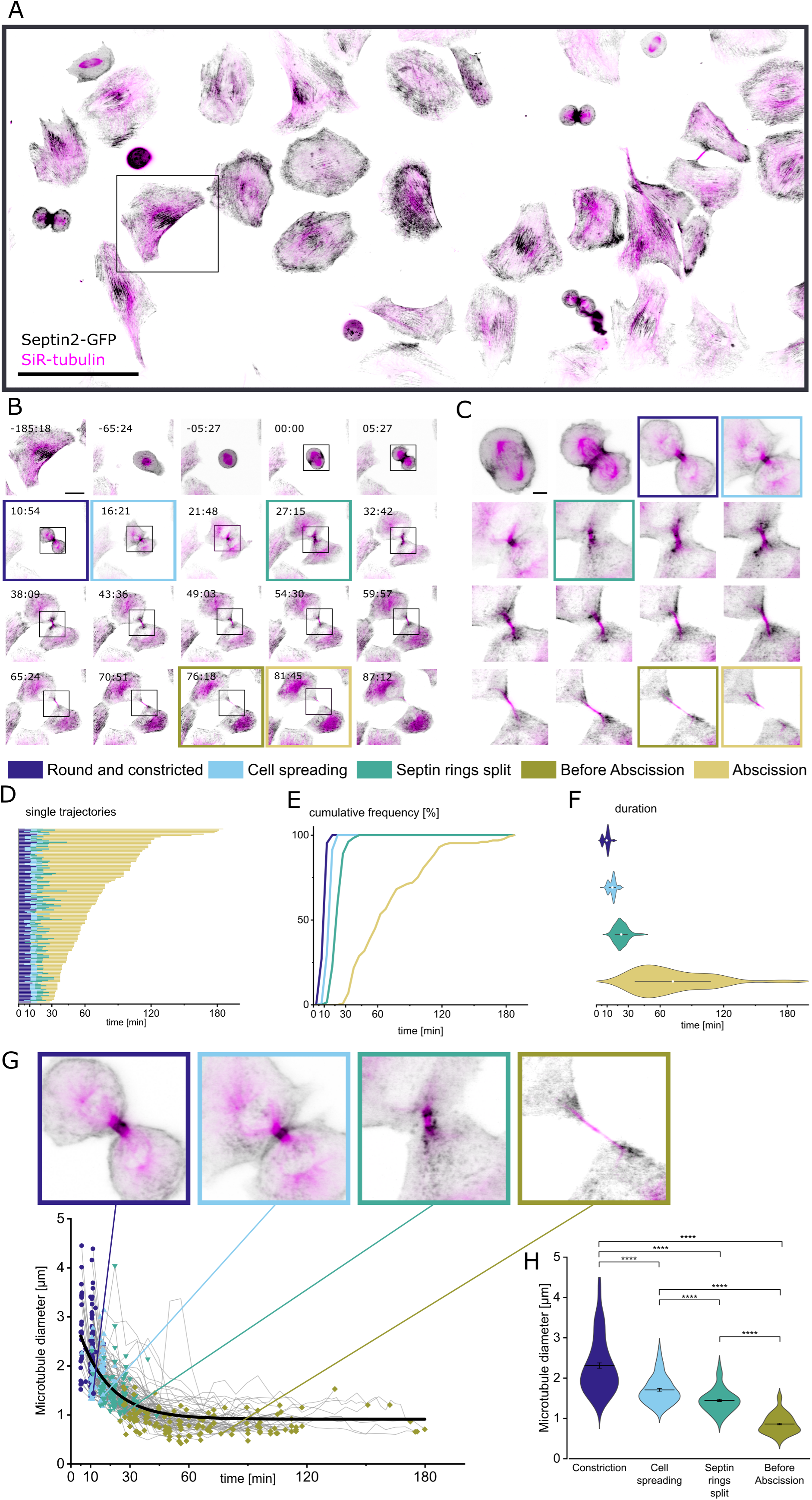
Quantitative lattice light-sheet live-cell imaging allows for characterization of cytokinetic progression based on changes in Septin2-eGFP ^EN/EN^ organization and microtubule-bundle thinning. A) Maximum intensity projection of the first frame of a lattice light-sheet movie. Synchronized NRK49F cells expressing Septin2-eGFP ^EN/EN^ (inverted grey) were stained with SiR-tubulin (magenta) and imaged every 5:27 min for 10 hours. Scale bar is 100 µm. B) Cytokinetic trajectory of the single cell marked in (A) with characteristic events highlighted. Dark blue: Full constriction of the cleavage furrow, light blue: spreading of the daughter cells on the substrate, turquoise: splitting of the tubular septin structure into septin rings, olive: the last frame before abscission, sand: abscission of the microtubule bundle in the ICB. Scale bar is 20 µm C) Zoom of the cell shown in (B). Scale bar is 5 µm. D) Plot of 129 single cell trajectories over the specific events (from N=7 movies). The start point t=0 is defined by the appearance of Septin2-eGFP ^EN/EN^ at the cleavage furrow. E) Cumulative frequency plot demonstrating the time passed until all cells reached a specific event. Shown are the same cells as in (D). F) Average duration between t=0 (appearance of Septin2-eGFP ^EN/EN^ at the cleavage furrow) and the respective event for all cells in (D, E). Events are: Round and constricted (“RC”, dark blue), Cell Spreading (“CS”, light blue), Septin Ring Split (“RS”, turquois), Abscission (“A”, sand). White circle with whiskers depicts the mean ± standard deviation. G) Plot of trajectories of measured microtubule-bundle diameter in the ICB over time for each cell. The cytokinetic events Constriction, Cell Spreading, Septin Rings Split and the last frame Before Abscission of the microtubule bundle are color-coded for all cells. Arrows point to the values measured for the cell depicted in (B) and (C). n=127 cells. The black solid line represents a single exponential decay fit with y=A*exp(-x/t)+ Y, Y = 0.915±0.017, A=2.32 ±0.09, t=15.8 ±0.8. R^2^=0.548 H) Shows the average microtubule diameter for each of the four cytokinetic events. Outliers were identified with the Grubbs test and masked. Mean ± SEM are shown. For statistical analysis, a Kruskal-Wallis test with post-hoc Dunn’s test was used. Means differ significantly amongst all groups. **** p > 0.001

We thus decided to use tubulin and Septin2-GFP^EN/EN^ as markers for classification in ultrastructure Expansion Microscopy^47^ (U-ExM, Figure 2A). Indeed, when we imaged tubulin in U-ExM, we found that the higher resolution and decrowding of the midbody revealed much more detailed features like the intersecting microtubule bundles at the center of the ICB, an area that in regular immunofluorescence microscopy remains unstained due to steric hindrance in antibody staining (Figure 2B, for quality control see Supplementary Figures S1-S3). We performed tubulin and Septin2-GFP^EN/EN^ staining in combination with specific staining of individual cytokinetic proteins in a third channel. We proceeded to stain for 18 proteins, DNA and a membrane marker and took a total of ∼ 2000 images in 20 datasets (Figure 2C).

**Figure 2:**
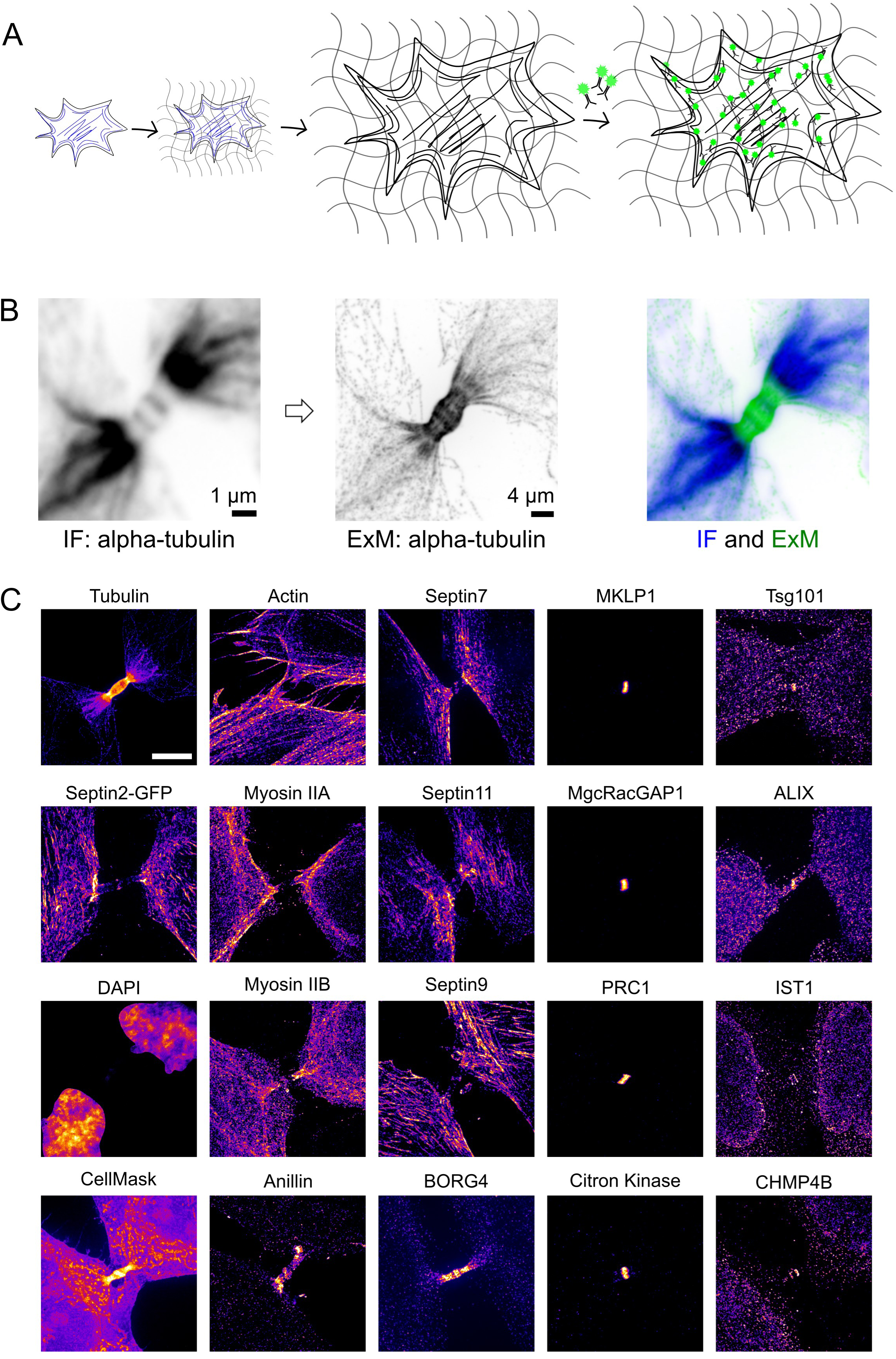
Expansion microscopy reveals the nanoscale organization of cytokinetic proteins at the midbody. A) Schematic illustration of the post-expansion staining process. Step 1: An immunostained cell (blue) is embedded into a hydrogel. Staining pre-expansion is optional and can be used to compare pre-and post-expansion staining quality as done in (B). Step 2: The hydrogel is homogenized and expanded. Step 3: In the expanded hydrogel, the protein of interest is probed again with an immunofluorescence staining (green). B) Left: Maximum intensity projection of a midbody stained with antibodies against alpha-Tubulin in regular immunofluorescence (IF). Middle: same cell after expansion and post-expansion staining of alpha-Tubulin. Right: Overlay of the midbody immunostained before (blue) and after gel expansion (green). C) Overview of all targets that were visualized in the ICB in this study. Representative single U-ExM images of the 20 targets as maximum intensity projections are shown. Images are individually adjusted for brightness and contrast and pseudo-colored by intensity with fire LUT. Scale bar is 5 µm.

The combination of the Septin2-GFP^EN/EN^ and the tubulin staining allowed the recognition of the defined stages as characterized in the lattice-light sheet data (Figure 3A). Importantly, the higher resolution allowed us to identify a population of dividing cells that showed a septin structure in the middle of the ICB and septin “manchettes” at the bridge-cell junction (Figure 3A, “septin manchettes”, SM). Characterizing this population as a cytokinetic stage was also supported by another observation. When we fixed cells at different time points after release from synchronization, we found that the newly described stage fit well into the progression of cytokinetic stages from round and constricted (RC) cells to full abscission (A) (Figure 3B).

**Figure 3:**
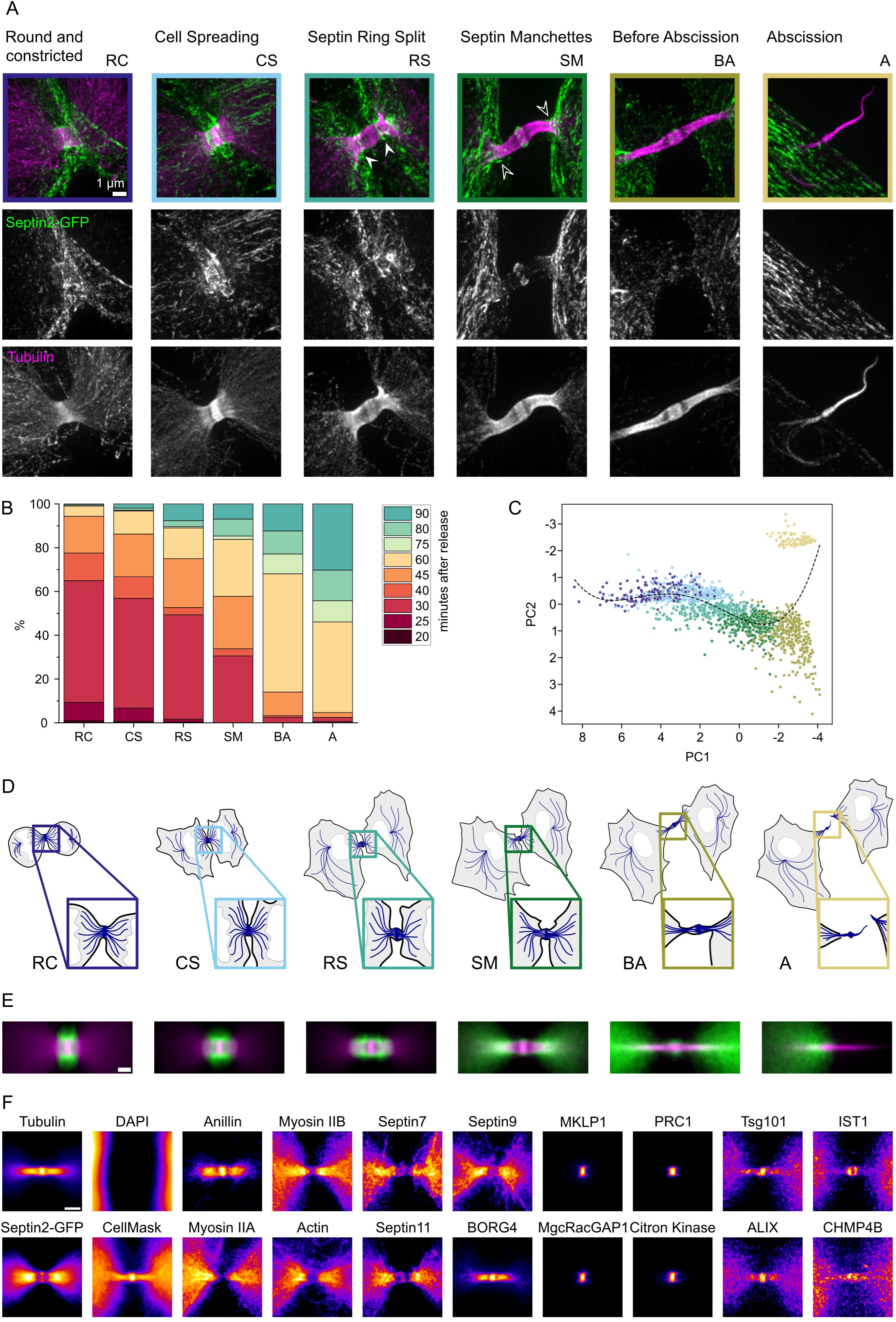
Expansion microscopy reveals six distinct stages in cytokinesis including the “Septin Manchettes” as an additional step before abscission. A) Selection of Septin2-GFP and tubulin-stained U-ExM images from 2068 individual cells representing six recognized stages in cytokinesis: Round and constricted (“RC”, dark blue), Cell Spreading (“CS”, light blue), Septin Ring Split (“RS”, turquois), Septin Manchettes (“SM”, dark green), Before Abscission (“BA”, olive), Abscission (“A”, sand). Septin2-GFP^EN/EN^ (green) and tubulin (magenta) are shown in merges and as individual grayscale maximum intensity projections. Solid white arrowheads point towards the split septin rings, outlined white arrowheads point to the septin manchettes. Note that the Septin Manchettes stage could only be identified in the high-resolution U-ExM dataset, not in the LLS movies (compare Figure 1). Scale bar is 1 µm. B) Histogram showing the correlation between the time of fixation after release from cell cycle synchronization and the appearance of the six cytokinetic stages. A total of 1930 images of ICBs was analysed. C) Principal component analysis of the U-ExM images used in this study. A total of 1696 images of ICBs was analysed. The polynomial fit (black dotted line) of the data represents the pseudotime axis. D) Schematics of the identified cytokinetic stages. Note the addition of the Septin Manchettes (SM) stage, which could only be identified in the high-resolution U-ExM dataset. E) Averaged mean intensity projections of the U-ExM dataset. Septin2-GFP (green) and tubulin (magenta) are shown. Scale bar: 1 µm. The following number of images was used to generate the averages: RC: n=222, CS: n=320, RS: n=312, SM: n=345, BA: n=405, A: n=326. F) Averaged mean intensity projections of the 20 targets that were visualized for this study in the *Septin Manchettes* stage. Scale bar is 2 µm. The following number of images was used to generate the averages: Tubulin n=345, DAPI n=345, Septin2-GFP n=345, CellMask n=21, Anillin n=18, Myosin IIA n=16, Myosin IIB n=26, Actin n=27, Septin7 n=12, Septin11 n=15, Septin9 n=15, BORG4 n=14, MKLP1 n=12, MgcRacGAP1 n=18, PRC1 n=16, Citron Kinase n=18, Tsg101 n=20, ALIX n=18, IST1 n=35, CHMP4B n=13. Images were individually adjusted for brightness and contrast and pseudo-colored by intensity with the fire LUT.

To ask if an identifiable pattern was underlying our observations, we used a combination of manually and automatically extracted, object-based features from Septin2-GFP^EN/EN^ and tubulin-stained cells in our dataset to perform a principle component analysis (PCA, Figure 3C). We selected these features based on their ability to describe the changes in shape and size of the microtubule bundle in the ICB and the cell body and the distribution of Septin2-GFP ^EN/EN^ along the ICB. The contribution of each of these features to PC1 and PC2 is summarized in Table 1. Especially the size of the overall spindle and ICB area and the spread of the septin structures along the ICB contributed to the variance of the data. PCA of these features showed all stages as individual clusters and especially the new “septin manchettes” stage, whilst remaining a distinct population, aligned well between the earlier “septin-ring split” and the later “before abscission” stages (Figure 3C). Our system could thus distinguish even stages that differed in small detail very well and assign them on a pseudotime axis. Only the spreading of previously rounded cells on the substrate after cleavage-furrow ingression was not distinguished very clearly, likely because the ExM images never showed entire cells due to their expanded size. We next aligned all images in the Septin2-GFP^EN/EN^ and tubulin channels for the specific stages to generate an average nanoscale 3D map of the ICB over time (Figure 3D and E). Importantly, even though hundreds of images were used for each averaged image, the stages showed distinct signatures in septin and tubulin staining, confirming that the observed patterns were indeed present in the said stages and not evened out by averaging (Figure 3E). We then averaged the images of all markers for all stages and found that many of them showed stage-specific staining as well, consistent with function-related reorganisation (see Figure 3F for averaged images of all markers in the “septin manchettes” or “SM” stage).

**Table 1:**
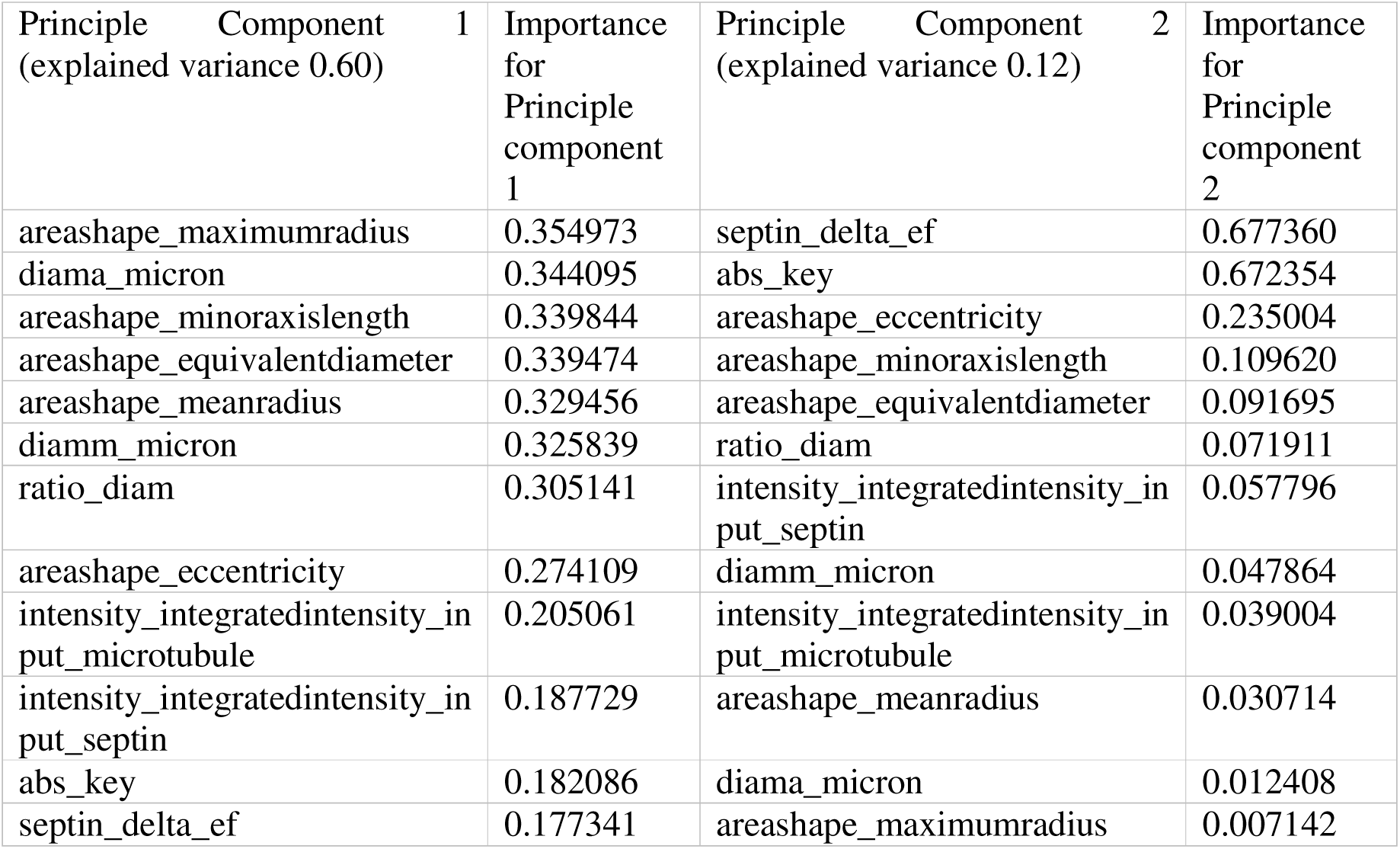
Ranked list of features and their contribution to PC1 and PC2 in PCA analysis of cytokinesis.

### Nanoscale analysis of cytokinetic proteins over time

Using our pseudotime, color and 3D analysis, we assembled a 5D dataset of cytokinesis. This dataset contains the averaged superresolved images of 18 proteins, the plasma membrane and the nucleus organized into the 6 stages and is navigable in the three spatial dimensions. This dataset allowed us to ask, how any protein was organized at the nanoscale in respect to any other of the proteins we measured, and is accessible via https://cytokinesismap.org (See also Supplementary Files 1-3). The experimental pipeline is described in Supplementary Figure S4. In the following, we describe several observations our dataset allowed us to make that have previously not been reported.

### Actomyosin

The constriction of the cleavage furrow and further midbody thinning are mediated by a contractile actomyosin network at the plasma membrane (Figure 4A). Consistent with this, when we averaged hundreds of U-ExM images of actin, non-muscle myosin isoforms IIA (NM IIA) and IIB (NM II2B) and Septin2-GFP^EN/EN^, we found membrane-apposed myosin IIA & IIB structures in early cytokinetic stages (Figure 4B). Despite their common localization at the plasma membrane during constriction, we could observe clear differences in organization between actin, myosin IIA & IIB and Septin2. Septin2 staining was consistently found outside of detected myosins, suggesting a more directly membrane-apposed organization of septins (Figure 4B, D, E). As the dynamic ICB exhibited a number of filopodia and small membrane protrusions (see also Figure 4A), the averaged actin staining that likely represented the membrane cortex was found in a deep layer also outside of septin staining in the averaged images in the early stages (Figure 4B, E). This is consistent with the extensive membrane folding and filopodia that are observed at the ICB as the cell reorganizes the plasma membrane material pulled together in cleavage-furrow ingression. Furthermore, myosin IIA was consistently found to the “cellular side” of myosin IIB, and myosin split into separate rings well before septin-staining did (Figure 4B, C). As the cells spread, the myosin staining accumulated in two nicely separated disks spaced about 1 µm apart that penetrated the microtubule bundle, while Septin2-GFP^EN/EN^ and actin remained peripheral (Figure 4D-F) and continued to stain the entire ICB membrane in a tube-like fashion (Figure 4B). Strikingly, while myosins never became visible in the midbody, Septin2-GFP^EN/EN^ showed a distinct central midbody staining until the “septin manchettes” stage (Figure 4B, C). As myosin disks split further apart, Septin2-GFP^EN/EN^ staining was enriched in rings surrounding the myosin disks while assuming a globular diffuse plasma membrane localization at the bridge as well as the cell bodies (Figure 4D-F). As the septin manchettes formed at the hillock of the ICB, myosin staining still filled the outer arms of the ICB, supported by cortical actin (Figure 4B, E). As the ICB constricted further towards abscission, the myosin disks similarly reduced in diameter and disappeared from the arms of the bridge as actin remained on the cortex (Figure 4E, F). The clear layering (from membrane to center) Septin-actomyosin-Myosin IIA&B/Tubulin at the “septin manchettes” stage is apparent in the midbody cross-section (Figure 4E, F) and hints at the transmission of actomyosin contractile forces towards the plasma membrane through septins to support secondary ingression of the ICB.

**Figure 4:**
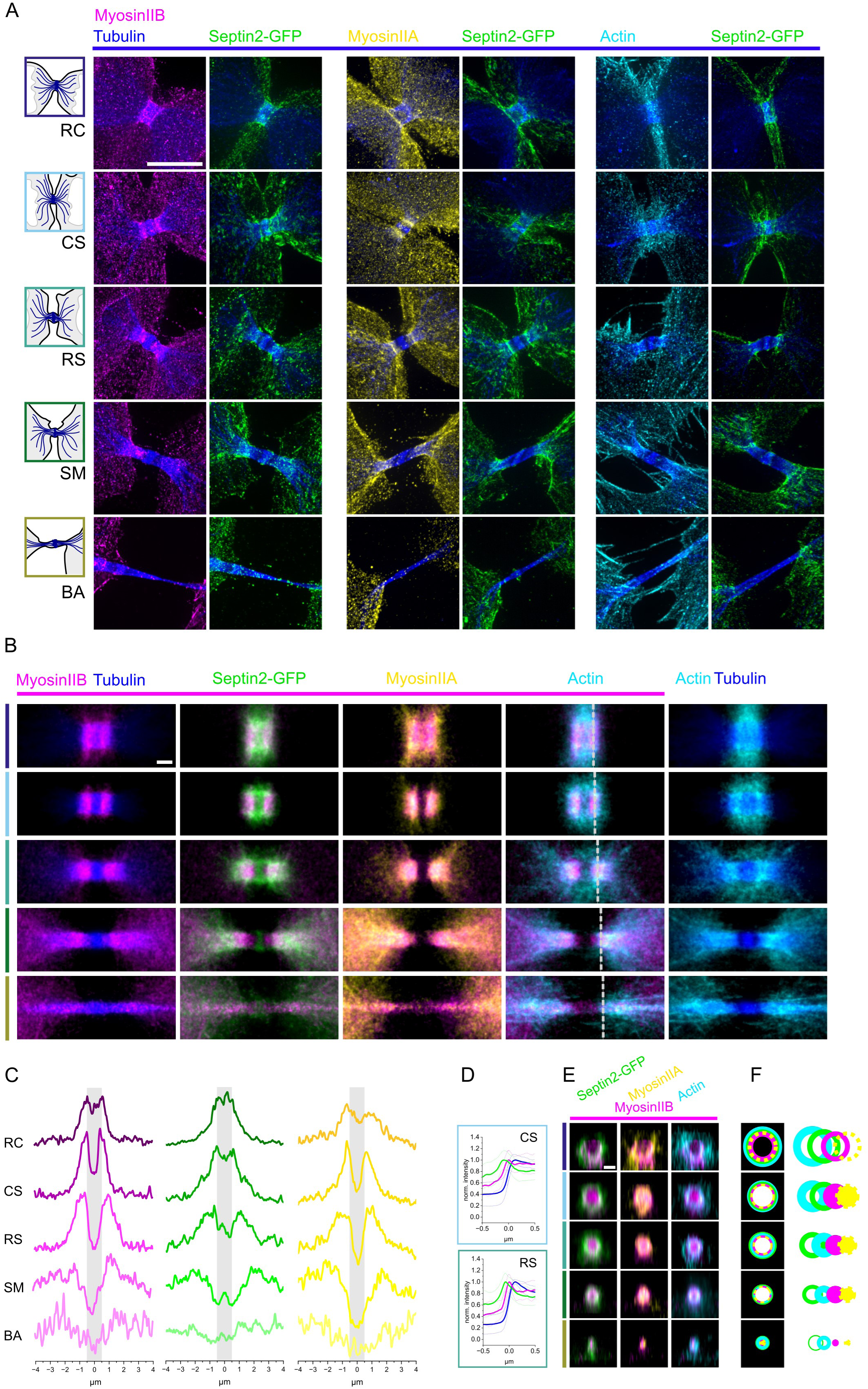
Myosin rings split at secondary ICB ingression before septin ring splitting, and myosins are located to the cytosolic side of septin staining. A) Representative U-ExM images of five different cytokinetic stages. Cells were immuno-stained with against myosin IIB (magenta), alpha-Tubulin (blue) and Septin2-GFP (green), myosin IIA (yellow) and beta-Actin (cyan). Maximum intensity projections are shown. Scale bar is 5 µm. B) Pseudocolor merges of averaged mean intensity projections of myosin IIB staining (magenta) with alpha-Tubulin (blue), Septin2-GFP (green) myosin IIA (yellow), actin (cyan) and based on the stages depicted in (A, top to bottom). Rightmost: Pseudocolor merge of averaged mean intensity projection of actin (cyan) and tubulin (blue). Scale bar is 1 µm. C) Line profiles of mean intensity along the division axis for each stage to emphasize the change in relative distribution of myosin IIB (magenta), Septin2-GFP (green) and myosin IIA (yellow) over pseudotime. D) Normalized and averaged line profiles of Septin2-GFP (green) and Myosin IIB (magenta) across the ICB arm in the Cell Spreading (“CS”) and Ring Splitting (“RS”) stage. The stained central microtubule bundle (blue curve, positive x-values correspond to the area of the microtubule bundle) indicates that Myosin and septin form layers between the microtubule bundle and the plasma membrane. Dotted lines represent the standard deviation. E) Orthogonal sections across the diameter of the 3D-averaged ICB as indicated by the grey line in (B) shows the radial distribution of Septin2-GFP, Myosin IIA, Myosin IIB and actin. Scale bar is 1 µm. F) Schematic illustration of change in distribution of the acto-myosin machinery across the ICB diameter.

### Septin and Anillin

Anillin is an essential, conserved adaptor protein in cytokinesis that binds actin and regulatory molecules and recruits septins to the plasma membrane^41,50^. However, it is not clear, how its roles in signaling and cytoskeletal regulation are executed over time in cytokinetic progression. Using our assay, we dissected the relative distribution of Anillin and septin over time in the evolving midbody. In single U-ExM images, it appeared that after initially strong colocalization with Septin2-GFP^EN/EN^, Anillin at later stages assumed a more microtubule-focused organization (Figure 5A). After cleavage-furrow constriction, Septin2-GFP^EN/EN^ and Anillin colocalized completely in a ring that lined the almost perfectly circular connecting tube between the daughter cells and this persisted during cell spreading. However, beginning at this stage, Anillin staining started to appear in the center of the midbody, on the microtubule bundle. This marked the appearance of two distinct pools of Anillin, one colocalizing with Septin2-GFP^EN/EN^ at the periphery and one accumulating on the microtubule bundle (Figure 5B, C). This trend of Anillin moving from membrane to midbody persisted until abscission, when Septin2-GFP^EN/EN^ at the midbody membrane was almost completely lost and Anillin colocalized with the, now thinner, midbody microtubule bundle. It seemed as if Septin2-GFP^EN/EN^ was required for Anillin membrane localization and that in the absence of Septin2-GFP^EN/EN^, Anillin would preferentially bind to microtubules as especially apparent in midbody cross-sections (Figure 5D, E). Indeed, midbody-associated Anillin staining further constricted with microtubule bundles until abscission. In contrast, the Anillin-interacting, persistently midbody-localized MgcRacGAP1 filled the midbody volume at the center, visibly extending beyond and surrounding tubulin (and Anillin) staining (Figure 5F, G). This nanoscale compartmentation of molecules within the central midbody domain lead us to ask, whether we could detect a further specialized localization of midbody molecules.

**Figure 5:**
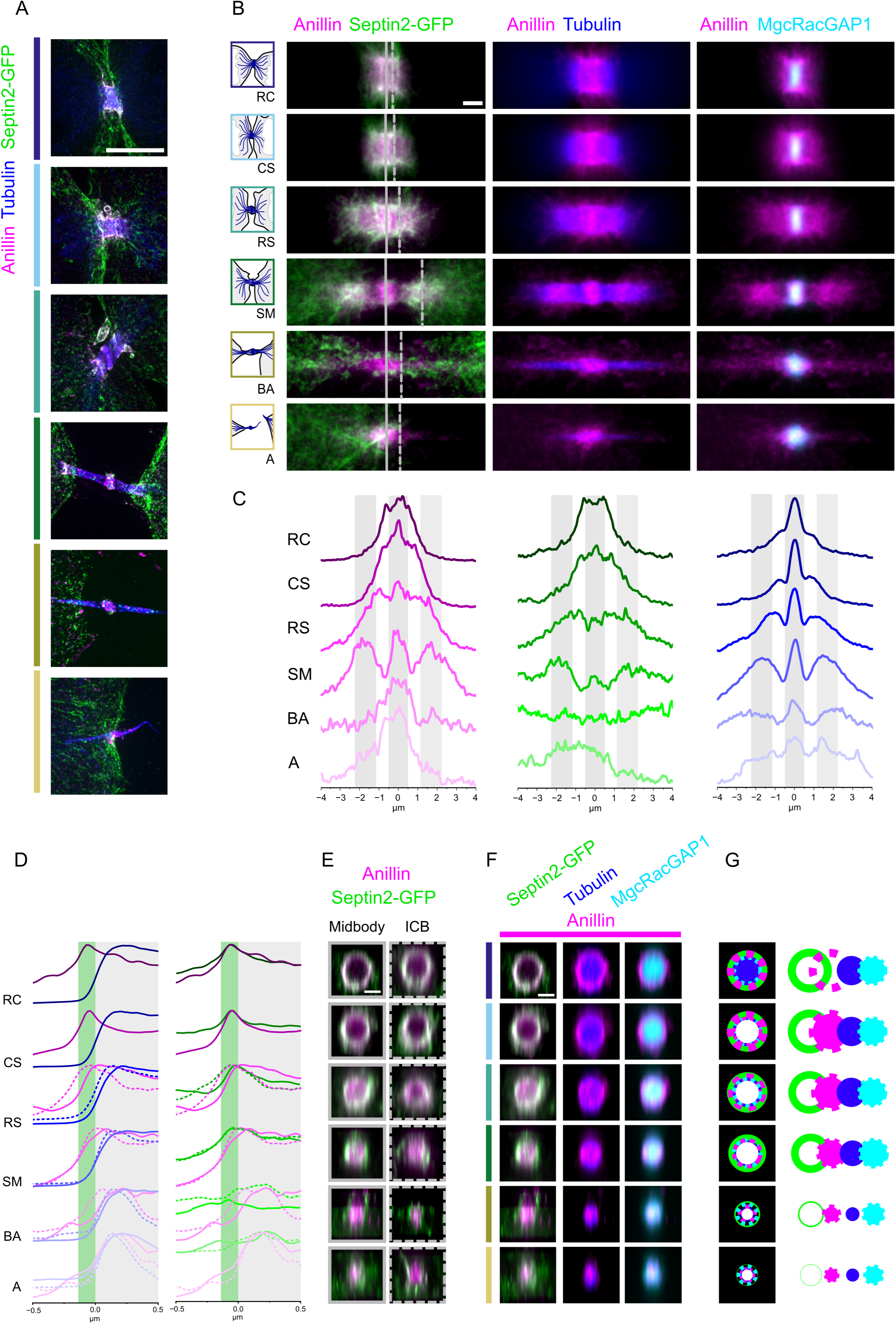
Anillin shifts from a membranous, Septin-associated state to microtubule-bundle and midbody association as abscission approaches. A) Representative U-ExM images of the six different cytokinetic stages. Cells were immuno-stained against Anillin (magenta), alpha-Tubulin (blue) and Septin2-GFP (green). Maximum intensity projections are shown. Scale bar is 5 µm. B) Averaged mean U-ExM pseudolored intensity projections of Anillin (magenta), tubulin (blue), Septin2-GFP (green) and MgcRacGAP1 (cyan) based on the stages depicted in (A). Solid and dotted lines indicate the position of orthogonal sections as shown in (E) and (F). Scale bar is 1 µm C) Line profiles along the division axis of individual cells were normalized and averaged per stage to depict the distribution of Anillin (magenta), Septin2-GFP (green) and tubulin (blue) over pseudotime. D) Line profiles across the edges of the ICB arm (dotted line) and the midbody (solid line) of individual cells were normalized and averaged per stage to depict the respective distribution of Anillin (magenta), Septin2-GFP (green) and tubulin (blue) over pseudotime. Green and grey bars mark the peripheral peak of Septin2-GFP staining and the microtubule bundle, respectively. E) Orthogonal sections across either the midbody (solid line) or the ICB arm (dotted line) in 3D-averaged volumes depicting the distribution of the Anillin (magenta) and Septin2-GFP over pseudotime. Scale bar is 1 µm. F) Orthogonal sections across the midbody in 3D-averaged volumes show the respective localization of Septin2-GFP, Anillin, tubulin and MgcRacGAP1. Scale bar is 1 µm. G) Schematic illustration of the distribution of Anillin in the midbody with respect to the microtubule bundle, Septin2-GFP and MgcRacGAP1.

### Nanoscale midbody compartmentation

To investigate nanoscale midbody compartmentation, we compared staining of PRC1, CIT (Citron kinase) and centralspindlin complex proteins MgcRacGAP1 and MKLP1. All these molecules localized to the central midbody throughout cytokinesis. However, when we compared the average 3D distribution of these molecules over time, we found strikingly different patterns (Figure 6A). The components of midbody organizing centralspindlin complex MgcRacGAP1 and MKLP1 first localize tightly to the intersecting microtubules in a homogenous, disc-like staining, that later widened over the edge of the microtubule cross-section in a developing midbody bulge after constriction and septin ring split (Figure 6A).

**Figure 6:**
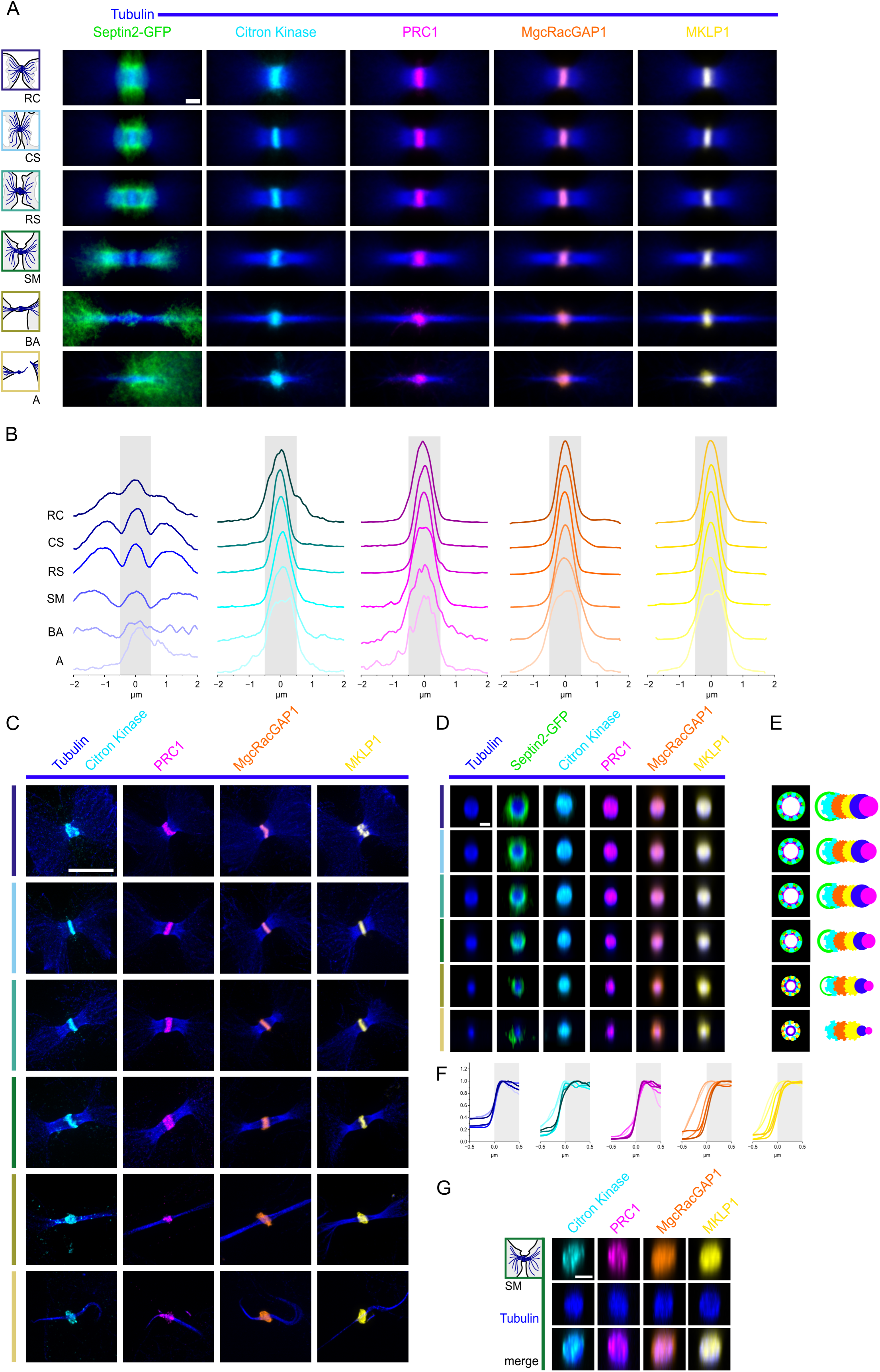
Organization of midbody-associated proteins in pseudotime. A) Averaged, pseudocolored mean intensity projections of tubulin (blue), Septin2-GFP (green), Citron Kinase (cyan), PRC1 (magenta), MgcRacGAP1 (orange) and MKLP1 (yellow) based on six cytokinetic stages. Scale bar is 1 µm. B) Profile lines along the division axis of individual cells were normalized and averaged per stage to emphasize the redistribution of tubulin (blue), Citron Kinase (cyan), PRC1 (magenta), MgcRacGAP1 (orange) and MKLP1 (yellow) over pseudotime. C) Representative single U-ExM images of each marker for the six different cytokinetic stages as depicted in (A). Maximum intensity projections are shown. Scale bar is 5 µm. D) Orthogonal sections in 3D-averaged volumes show the distribution of Tubulin (blue), Septin2-GFP (green), Citron Kinase (cyan), PRC1 (magenta), MgcRacGAP1 (orange) and MKLP1 (yellow) across the midbody over pseudotime. Scale bar is 1 µm. E) Schematic illustration of the redistribution of the midbody-associated proteins with respect to Septin2-GFP and the microtubule bundle. F) Normalized and averaged line profiles across the midbody edge are shown for tubulin (blue), Citron Kinase (cyan), PRC1 (magenta), MgcRacGAP1 (orange) and MKLP1 (yellow). Lighter colors indicate later cytokinetic stages (as in B). The grey bar marks the central microtubule bundle. G) Orthogonal sections of individual midbodies in the Septin Manchettes stage show the distribution of Citron Kinase, PRC1, MgcRacGAP1 and MKLP1. Note the punctate, discreet staining of Citron Kinase and PRC1 in contrast to the homogenous distribution of MgcRacGAP1 and MKLP1. Scale bar is 1 µm.

Occasionally, we observed it in septin-positive membrane outbursts that emanated from the midbody bulge. As the microtubule bundle thinned towards abscission, centralspindlin staining spread bi-directionally along the bundled microtubules as well (Figure 6B).

Citron kinase on the other hand shifted from the plasma membrane to the midbody during constriction. In contrast to centralspindlin, it exhibited a more punctate organization throughout cytokinesis. This we observed both at the midbody as well as in occasional spots that decorated the membrane after cell spreading, consistent with its role in myosin regulation (Figure 6C, G). Overall, the diameter of citron kinase staining followed centralspindlin at the midbody (Figure 6D-F). As septin manchettes formed and the microtubule bundle thinned further, citron kinase spots decorated the surface of microtubule bundles emanating from the midbody (Figure 6C). These spots spread wider than centralspindlin towards abscission (Figure 6B).

Finally microtubule-bundling protein PRC1 remained attached to the microtubule bundle in a spotty pattern, consistent with the presence of clusters of PRC1 as they are observed when motor-driven antiparallel microtubule pulling experiences “breaking” events^51^. Along the ICB axis, we found PRC1 localization to follow centralspindlin until the septin rings split.

Afterwards, it appeared persistently wider along the microtubule bundle than Citron kinase or centralspindlin until abscission (Figure 6A-C), suggesting that intersecting microtubule ends in the midzone are destabilized. In contrast to the other midbody proteins, the longitudinal spread of PRC1 was confined by the diameter of the microtubule bundle (Figure 6D-F). The spotty pattern of PRC1 (as well as Citron kinase) in midbody cross-sections could be observed in averaged as well as individual midbodies (Figure 6D and G, respectively), meanwhile centralspindlin always formed an evenly stained disk. Taken together, our observations allow unprecedented insight into midbody organization during cytokinesis.

### ESCRT

Finally, we analyzed the distribution of ESCRT complex molecules on the midbody, as these molecules are essential for abscission. We stained expanded cells with antibodies against ESCRT-I protein Tsg101 and the ESCRT-III molecules IST1 and CHMP4B, respectively, to ask, whether their organization was temporally coordinated with other cytokinetic events. Tsg101 was the first to appear at the midbody as soon as cells started to spread (Figure 7A) and remained attached to the midbody until after abscission. However, while first it was tightly focused on the midbody, it later became localized more diffusely emanating from the midbody bulge. Similar staining was observed for ALIX (see Supplementary Files 1-3). On the other hand, both ESCRT-III proteins IST1 and CHMP4B were first localized to punctate structures in the cytosol until they appeared associated with the bridge from the “septin manchettes” (SM) stage onward (Figure 7B-D, Supplementary File 4). These molecules formed ring-like structures to the right and left of the midbody (Figure 7C and Supplementary Figure S5). When rings were present on both sides of the midbody (sub-stage 2) occasionally we would observe a cone on one side (sub-stage 3). Midbodies with cones on both sides of the midbody (sub-stage 4) were observed seldomly, suggesting that one cone was sufficient for scission of the microtubule bundle (Figure 7E).

**Figure 7:**
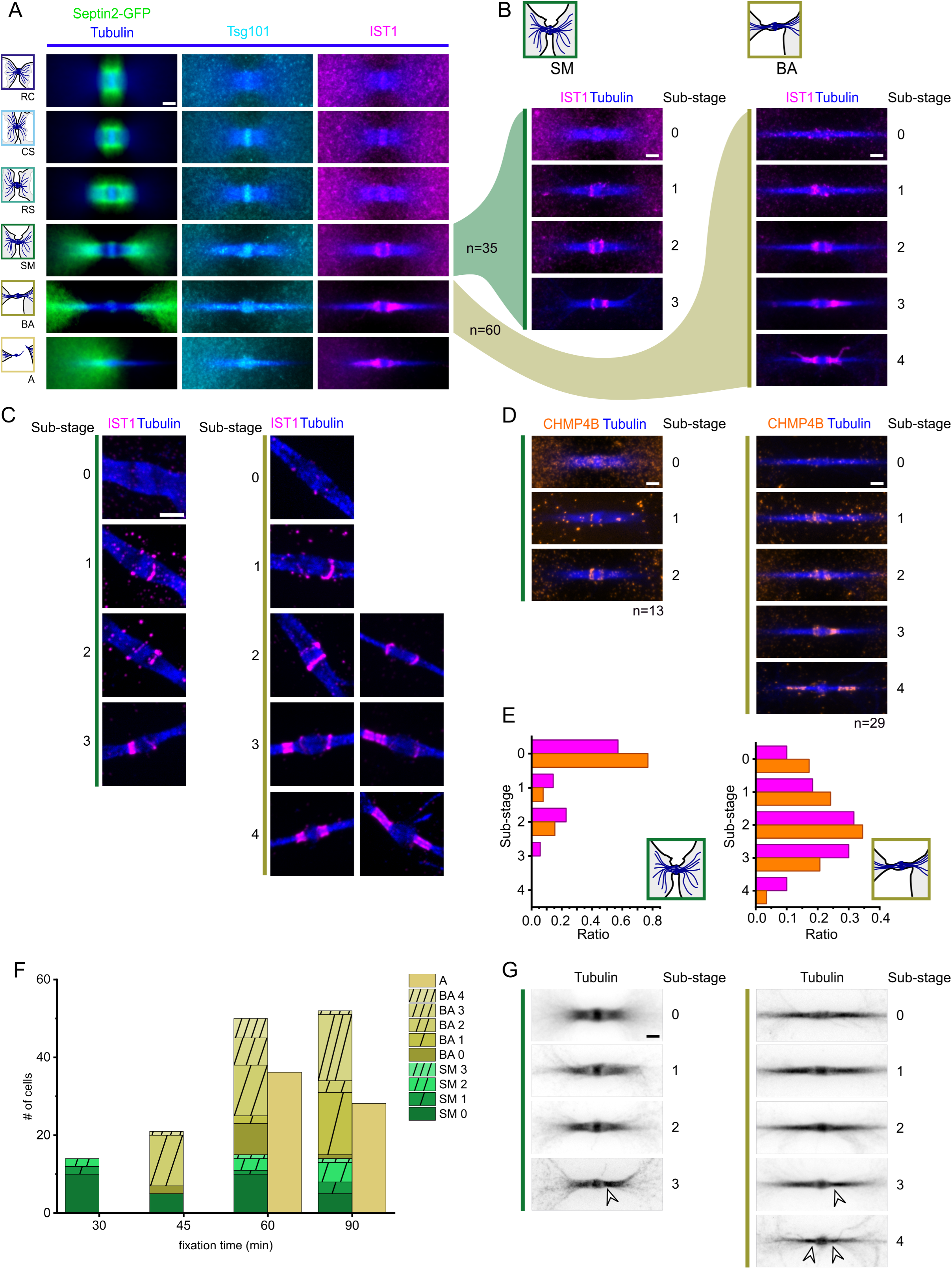
The spatial organization of ESCRT-III before abscission. A) Averaged pseudocolored mean intensity projections of tubulin (blue), Septin2-GFP (green), Tsg101 (cyan) and IST1 (magenta) based on six cytokinetic stages. Scale bar is 1 µm. B) Sub-stages of the IST1 organization at the ICB throughout the *Septin Manchettes* and *Before Abscission* stages. Averaged mean intensity projections of IST1 and Tubulin. Scale bar is 1 µm. C) Representative single U-ExM images of IST1 and Tubulin depicting the sub-stages. Scale bar is 1 µm. D) Similar sub-stages as in (B) could be identified in averaged mean intensity projections of CHMP4B (orange). Scale bar is 1 µm. E) Histograms of the ESCRT-III sub-stages show equal distribution for the targets IST1 (magenta) and CHMP4B (orange). F) Histogram of ESCRT-III sub-stages in correlation to fixation time. G) Averaged mean intensity projections of Tubulin in the sub-stages as shown in (B) and (D). White arrowheads point to constricted microtubule-bundles at sites of ESCRT-III spiral positions in (B and D).

The assembly of ESCRT-III rings and cones at the midbody was not correlated with any detectable changes in the pattern of septin localization. In effect, it seemed that ESCRT-III could assemble once septins were replaced form the midbody membrane. The appearance of ESCRT-III proteins in the early “septin manchettes” stage could mean that maturation from SM to “before abscission” (BA) stage happens independently from ESCRT-III assembly. This notion was supported by the observation that when we fixed cells 30 minutes after release from synchronization we observed only cells in the SM stage with ESCRT-III structures (about 28%, see Figure 7F), but no cells in the BA stage. When cells were fixed in another experiment 45 minutes after release, we already found the majority to be ESCRT-III positive, however, all of these were in the BA stage. Cells with completed abscission were first observed in the 60 min pool, together with cells in the SM and BA stages with many stages of ESCRT-III assembly. The ESCRT-III positive share rose again in the 90 min fixation timepoint, where we found both SM and BA stage cells to be overwhelmingly ESCRT-III positive. This suggested that ICBs can develop from the SM to the BA stage independently of ESCRT-III assembly but not abscise (we never found cut bridges without ESCRT-III). At the 60 minutes timepoint, more than 50% of the bridges had evolved to before abscission with two or more ESCRT-III rings and abscission was detected as well. In the pool fixed 90 minutes after release, we found the largest population of before abscission bridges with asymmetrical ESCRT-III structures, especially the sub-stage with one ring is very prominent. This hints at the parallel assembly of an ESCRT-III double ring on either side of the midbody as a fast, abscission rate controlling step. Strikingly, almost all cells showed some form of ESCRT-III staining 90 min after release, suggesting that in the SM stage ESCRT-III can mature before the BA stage is reached. On the other hand, the presence of ESCRT-III negative BA stage cells demonstrated that cells can move to the BA stage without having ESCRT-III. This strongly suggests that ESCRT-III recruitment and maturation of the ICB are not interdependent processes, but may occur separately from each other. The developing ESCRT-III cones and spirals further narrow the bridge microtubules, which is reflected in indentations in the microtubule bundles at the sites of the cones (Figure 7G).

### Pseudotime visualization of the intercellular bridge

Finally, we associated the principal component data point for every image onto its closest point on the fit pseudotime function to create a 1D pseudotime axis (Figure 3C). Sampling the images along the pseudotime axis thus represents progression through cytokinesis.

Averaging several images in a moving window allowed us to generate a video of the spatiotemporal development of Septin2-GFP ^EN/EN^ and tubulin in the intercellular bridge (Supplementary Movie 2). Using an average of 98 images that progressed for 49 images with each frame resulted in a series of 34 frames played at 10 fps. This video clearly demonstrated that the transitions between the 6 classified stages happened in a gradual but continuous way, consistent with the claim that our pseudotime analysis recapitulates the natural progression of cytokinesis over time. When we included MgcRacGAP1 and IST1 in this analysis, we likewise could visualize a continuous change over time (Supplementary Movie 3).

MgcRacGAP1, as expected appeared as a brightly stained, homogenous disc at the midbody early on and later seemed localized to a larger bulbous membrane structure with an often-shifting appearance due to lower averaging in the video. This is consistent with our observation that membranous outbursts of MgcRacGAP1from the midbody bulge were observed from the SM stage onward and that averaged MgcRacGAP1 staining extended beyond the midzone microtubule bundle as a result. Visualization of ESCRT-III assembly (IST1) in this video exhibited nearly continuous accumulation to cones just before abscission, further supporting the notion that our pseudotime analysis represents the natural progression timeline. When we plotted the stage (SM or BA) and ESCRT-III assembly state (absent to double cone, from 0-4 as in Figure 7) against pseudotime, we clearly found progression from low ESCRT-III assembly to high ESCRT-III assembly states (see Supplementary Figure S6). Strikingly though, we also found that there was a clear distinction between SM and BA stages. Early stages were low ESCRT-III-SM, while high ESCRT-III-SM states overlapped with low ESCRT-III-BA states and high ESCRT-III-BA stages came last. These data again strongly support our conclusion that ICB development and ESCRT assembly are independent phenomena.

## Discussion

We here describe an assay for the superresolved multiplexed investigation of cytokinesis in animal cells by correlated live-cell and Expansion microscopy and pseudotime analysis.

Since current microscopy studies of cytokinesis are always limited to 2-3 color channels and relatively low temporal and spatial resolution, a comprehensive, dynamic picture of the relative organization of the essential components was inaccessible. Here we have imaged 20 targets over the entire process and generated a detailed multidimensional map of cytokinesis. Doing so, we generated an averaged, nanoscale map of the components that could resolve several decisive steps with previously inaccessible accuracy. In this way, our assay allows us to visualize the relative organization of cytokinetic molecules with high confidence and resolution. This resulted in the discovery of nanoscale reorganizations of the constriction machinery as well as midbody proteins and septo-anillin. We furthermore categorize cytokinesis into stages that are well defined by Septin2-GFP and tubulin staining and correlated with specific cellular functions. In fact, the features that allow the classification into stages also allow us to create a pseudotime progression of cytokinesis from all our images and a nanoscale movie of cytokinesis. A summary of our findings is illustrated in Figure 8, where we emphasize the specific reorganization events we could here resolve to the nanoscale in averaged images between the individual stages. Since it is not feasible to show channel merges of all markers, we also provide a schematic of the distribution of 10 representative markers for every stage from the perspective of the intercellular bridge and the midbody, respectively. This resource is publicly available under: https://cytokinesismap.org, where a navigable file of all 20 components in 3 dimensions over the 6 stages can be browsed (see also Supplementary Files 1-3).

**Figure 8:**
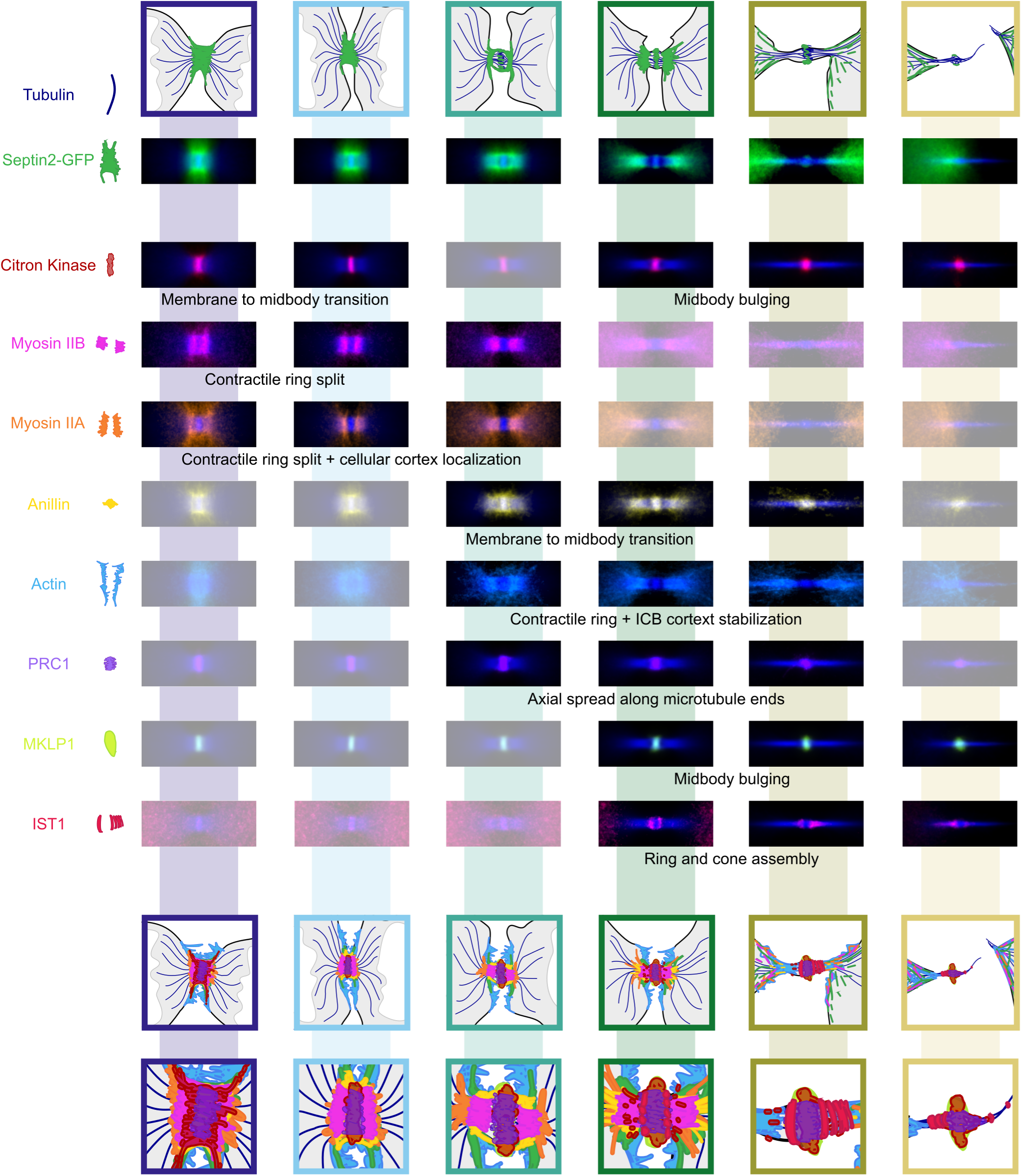
The development of the cytokinetic bridge over time in multiplexed superresolved pseudotime imaging. Shown are averaged mean intensity projections of the markers Septin2-GFP (green) and tubulin (blue) with the development of Citron Kinase (maroon), Myosin IIB (magenta), Myosin IIA (orange), Anillin (yellow), actin (light blue), PRC1(lavender), MKLP1 (lime, also representative of MgcRacGAP1) and IST1 (red, also representative of CHMP4B) throughout the six cytokinetic stages. Prominent reorganization events of molecules are highlighted. The cartoon in the bottom combines all depicted molecules and their relative organization at the ICB and in the midbody, respectively.

The feature-based classification of cell cycle microscopy images has been done previously by using chromatin morphology to classify mitotic stages using supervised machine-learning^52^ combined with a hidden Markov model to set constrains on the chronology of the analyzed time-lapse data. This however, is only possible when the previous and following steps of a snapshot image are known, i.e. in a continuous time-series. Multiplexed temporal imaging of cell division has also been performed using chromatin, cellular morphology and volume to model a mitotic standard time with 20 individuals steps and assign the localization of 28 proteins based on fluorescence confocal live-cell imaging^53^. This dynamic map however is limited in its resolution to the diffraction limit of light and stopped at constricted cells, excluding cytokinesis. However, it emphasizes the usefulness of symmetric processes such as chromatin separation during mitosis to sort events on a time-scale. For structures investigated by superresolution microscopy methods based on fixed cells as in our case, the combination of averaging and feature-based pseudotime extraction has for example been applied to nanoscale structures as the clathrin endocytic machinery in vesicle internalization^4,54^ and centriole development^5^. The features we employed here focused on the shape and size of the microtubule bundle in the ICB and the corresponding septin channel. Based on our LLS live-cell imaging, the notion of a thinning microtubule bundle over time was observed, as described by others^55^. However, this thinning showed high cell-to-cell variability, such that this single feature (diameter of the microtubule bundle) did not suffice to project a given snapshot onto a time-axis. By combining multiple bridge-related features, however, we could create a multidimensional feature-space that clearly differentiated between the 6 cytokinetic stages, sorted them in the expected chronology and allowed to visualize transitions between those stages. We then could average the relative distribution of several cytokinetic molecules over time and with this also their relative localization.

Averaging images can reveal general patterns of protein distributions with high confidence^56^ and convincingly visualize the assembly of symmetric multiprotein structures^57–62^. However, in large, asymmetric assemblies, information on non-homogenous distribution can be lost as we observed for the ESCRT-III spirals in Figure 7. It is thus important to be aware of such errors intrinsic to the technique when appreciating the averaged images. At the same time, when a structure is present in the averaged image, this is very strong evidence for it to reliably exist within a process. An example here are the indentations in microtubule bundles at sites of ESCRT-III recruitment in sub-stages 3 and 4 (see Figure 7). Likewise an offset between localizations of molecules is strong evidence for a layered organization of molecules even if it is only accessible in multiplexed images like we show for Myosin IIA and IIB or Septin2, Anillin, tubulin and MgcRacGAP1 respectively that were not imaged simultaneously, but only assembled as separate channels in the averaged multiplexed image.

We first defined stages arbitrarily from morphological features that reflected cellular processes in “round and constricted”, “cell spreading”,”septin ring split”, “before abscission” and “abscission”. (In this study, we have defined the severing of the microtubule bundle as “abscission”. However, we are aware that even after microtubules along the ICB are cut, the structure itself can persist much longer^27^.) The higher resolution expansion microscopy images allowed us to categorize more clearly the development in the late stages of cytokinesis between the septin ring split, where the compressed spindle emerges as an ICB and abscission, where it is cut. This remains a poorly defined period, where many events are coordinated to lead to final abscission, and the spatiotemporal organization remains unclear. Particularly, in our experiments, a distinct substep towards the classical image of a thin, elongated ICB with a bulbous midbody became apparent, which we called “septin manchettes”. Here, the split septin rings formed manchettes around the ICB where it emerged from the cell body and Myosin IIA and IIB on microtubules supported secondary ingression of the microtubule bundle. Midbody components still resided on the intersecting midzone microtubules and did not form a bulge around it yet. Anillin had shifted from the membrane to the microtubule bundle and actin had cleared the midbody, leaving a thin septin ring at the midbody membrane. In the PCA of all our images, this stage occupied a defined zone between the RS and BA stages that partially overlapped with either, suggesting that our stage had foundation in a biological transition. We thus suggest that with the “septin manchettes” stage we here describe a necessary organizational stage in cytokinesis. This is further strengthened by the observations that i) in the septin manchettes stage do we first detect ESCRT-III assembly, but never in previous stages. ii) early in the timecourse we observe only “septin manchette” cells with ESCRT-III, but no cells in the “before abscission” stage (Figure 7F) and iii) within the SM stage, ESCRT-III assembly progresses with pseudotime (Supplementary Figure S6). However, while our data suggest that ESCRT-III positive SM cells can mature into BA cells, we cannot rule out that abscission occurs occasionally already at the SM stage, especially as we could already observe indentation of bundled midbody microtubules at locations of ESCRT-III staining (see Figure 7G). We however think this is highly unlikely, as all cells we observe after abscission drastically deviate from the SM stage in microtubule bundle thickness, Anillin distribution, and the formation of a mature midbody bulge. It has recently been reported that caveolae at an “intercellular bridge (ICB)-cell interface”^21^ regulate membrane tension in cytokinesis and this ICB-cell interface may be the location of the septin manchettes. Our data suggest that ESCRT-III assembly might not be a reliable marker for progression of cytokinesis and that the septin manchettes stage is a necessary step in ICB maturation.

We find Myosin IIA on the cell-directed side of Myosin IIB similar to what has been observed in live-cell imaging before^63^. The tight association of myosin and actin on the arms of the ICB during RS and SM stages emphasizes their role in secondary ingression of the microtubule bundle^63^. However, with the higher resolution in space and time we could show that the site of secondary ingression is not automatically the site of abscission, as there is clearly a step in bridge maturation in which myosins leave the thin microtubule bundle before abscission and before the ESCRT-III cones have fully formed. Instead, we propose a third ingression of the microtubule bundle that happens independently of myosin-forces once ESCRT-III starts to assemble into cones. Of note, actin remained longer along the ICB arms than Myosin IIA and IIB, which is consistent with its recently described role in mediating ESCRT-dependent microtubule severing^27^.

Anillin has been observed to organize into three rings at the maturing ICB, of which the central ring surrounding the midbody was more stable compared to the dynamically exchanging Anillin double rings flanking the midbody^64^. This suggests that there are indeed different pools of Anillin at the ICB with different properties. We observed that Anillin on the midbody also penetrated the microtubule bundle after constriction of the cleavage furrow.

The shift of Anillin localization from the membrane-associated Septin2 to tubulin in the center of the midbody upon the split of the septin rings hints at the emergence of distinct Anillin pools that through their interaction with different partners^15,64–66^ might have distinct functions in cytokinesis.

Taken together, we envision our assay in combination with pharmacological and/or genetic treatments or optogenetic perturbations to allow for a detailed investigation of substeps of cytokinesis as we give access to the temporal domain in superresolution imaging at a scale compatible with averaging and statistical analysis. We here used the for ultrastructural preservation well-established U-ExM method^5,47^. Especially in combination with even higher expansion factors^67–72^ or Ex-STED^73^, this pipeline should allow for highest resolution of subcellular organization. Together with multiplexed imaging of dozens of molecules in large scale assemblies^74^ this will greatly contribute to our understanding of cytokinesis in particular and functional multiprotein assemblies in cells in general.

## Materials and Methods

### Cell culture and synchronization

Rat kidney fibroblast cells (NRK49F) with a homozygous insertion of eGFP at the Septin2 N-terminus^48^ were cultured in DMEM (Gibco, 31053028) supplemented with 4.5 g/l glucose, 10% v/v FCS (neolab, 2095ML500) and 1% v/v GlutaMAX (Gibco, 35050038). Cells were cultured at 37°C and 5% CO_2_ in a humidified incubator, and routinely tested for mycoplasma contamination. For synchronization, NRK49F Septin2-eGFP^EN/EN^ cells were seeded on Geltrex (Gibco, A1413302) coated coverslips. In brief, 18 mm glass coverslips were incubated with a 1:50 suspension of Geltrex in optiMEM (Gibco, 31985070) for 30 min at 37°C. Cells were seeded onto the coverslips and grown in full medium supplemented with 2 mM thymidine (Acros Organics, 226740050) for 22 h to stall the cells at S-phase. Following the thymidine block, the cells were washed four times for 2 min each with warm full medium and incubated for 3.5 h. To further increase cell cycle synchrony for fixed cell experiments, after thymidine treatment, mitotic spindle assembly was blocked by incubation with 20 ng/ml nocodazole (Acros Organics, 358240500) in full medium for 4 h. To release from the nocodazole block, the cells were carefully washed four times for 2 min each with warm full medium. Subsequently, cells started mitosis and were fixed at different time points during cytokinesis.

Synchronization and SiR-tubulin staining for live-cell lattice light-sheet imaging.

NRK49F Septin2-eGFP^EN/EN^ cells were seeded onto Geltrex-coated glass-bottom imaging dishes (Zellkontakt, 6160-30). They were synchronized by a single 2 mM thymidine block for 22 h and released into full medium containing 100 nM SiR-tubulin and 10 µM verapamil (both Spirochrome, SC002) and incubated for 5 h. Prior to live-cell imaging, medium was exchanged to FluoroBrite DMEM (Gibco, A1896701), supplemented with 100 nM SiR-tubulin and 10 µM verapamil.

Ultrastructure Expansion microscopy (U-ExM) of cytokinetic cells.

Cultured cells were fixed and processed using the U-ExM protocol^47^ with minor adjustments of the fixation and the gel homogenization. To obtain ICBs at different stages, cells were fixed at different time points after release from synchronization. The culture medium was removed and the cells were rinsed briefly with PHEM buffer (60 mM PIPES-KOH, 25 mM HEPES, 10 mM EGTA, 2 mM MgCl_2_, pH 6.9) warmed to 37°C. Cells were fixed in 4% v/v paraformaldehyde (PFA; Electron Microscopy Sciences, 15710)/0.1% v/v Triton-X 100 (VWR, 0694-1L) in PHEM buffer at 37°C for 15 min. Following fixation, the cells were rinsed with PHEM buffer and then placed in freshly made anchoring solution consisting of 0.7% v/v PFA and 1% v/v acrylamide (AA; Sigma, A4058-100ML) in PBS. The samples were incubated over night at RT.

U-ExM monomer solution (19% w/w sodium acrylate [Sigma Aldrich, 408220-100G], 10% w/v AA, 0.1% w/v N,N’-methylenbisacrylamide [Sigma, M1533-25ML] in PBS) was prepared at least one day prior to gelation and stored atc20°C. For gelation, 5 µl of 10% w/v ammonium persulfate (APS; Roth, 9592.2) and 5 µl of 10% v/v tetraethylenediamine (TEMED; Sigma, T9281-50ML) were mixed with 90 µl of monomer solution and briefly vortexed. The coverslips were removed from the anchoring solution, blotted dry from the side and placed on drops of 80 µl activated monomer solution on parafilm. They were incubated for 5 min on ice, and then placed at 37°C in a humid chamber for 1 h. Gels were transferred to wells of a 6-well plate containing denaturation buffer (200 mM sodium dodecyl sulfate [SDS; Sigma Aldrich, 75746-250G], 200 mM NaCl, 50 mM Tris, pH 9) and agitated for 15 min. Thereafter, the gels were separated from the coverslips and were placed in 15 ml centrifuge tubes filled with 4 ml denaturation buffer. The tubes were incubated at 95°C for 60 min. After denaturation, the gels were washed four times with MilliQ water and then placed in 10 cm cell culture dishes filled with 25 ml of MilliQ water. The gels were incubated for 30 min, after which the MilliQ water was replaced for a second 30 min incubation. Following a final water exchange, the gels were left to expand at 4°C over-night. The next day, the expansion factor of the gels was measured with a caliper and they were shrunk by two incubations in PBS lasting 15 min each. The shrunken gels were either processed for antibody staining or stored at 4°C for several weeks.

### Optimization of U-ExM protocol

To find the optimal conditions for continuous ICB staining, two aspects of the original U-ExM protocol were varied: The gels were subjected to increasing time of denaturation (15 min, 30 min, 45 min and 60 min) and different fixation agents were used upstream of the gel preparation. The different fixatives used were a) 4% v/v PFA/0.1% v/v Triton-X 100 in PHEM buffer, b) 4% v/v PFA in PHEM buffer, c) 4% v/v PFA/0.1% v/v glutaraldehyde (GA, Electron Microscopy Sciences, 16316) in PHEM buffer and d) methanol. For methanol fixation, after a rinse with PHEM buffer, the samples were incubated with ice-cold methanol for 20 min at-20°C. Subsequently, they were washed thrice for 10 min with PHEM buffer.

The PFA-based fixatives (a, b, c) were incubated with the samples for 15 min at 37°C, after which they were washed thrice with PHEM buffer for 10 min. Fixation was followed by anchoring, gel preparation and denaturation.

### Staining with CellMask

CellMask orange plasma-membrane stain (Thermo Fisher Scientific, C10045) was diluted 1:1000 in PHEM buffer. Synchronized and fixed cells were incubated on drops of CellMask solution for 1 h at RT. They were washed twice with PHEM buffer for 10 min each, followed by a post-fixation with 4% v/v PFA in PHEM buffer. The samples were washed again for 10 min with PHEM buffer, before being transferred to U-ExM anchoring solution.

### Staining of U-ExM gels

Squares of 1 cm side length were cut from the gels and transferred to a 24-well plate. Gels were incubated with primary antibodies, diluted in 2% v/v BSA/PBS at 37°C for 3 h with agitation. Next, they were placed in a 6-well plate and washed three times for 15 min with 0.1% Tween 20 (Roth, 9127.1) in PBS (PBS/T). Secondary antibodies and nuclear stains (See Tables 2-4 for a list of all staining used) were diluted in 2% v/v BSA/PBS and incubated with the gel pieces in a 24-well plate for 2.5 h at 37°C with agitation. Following the secondary antibody incubation, the gels were washed again three times for 15 min with PBS/T in a 6-well plate. For re-expansion, PBS/T was exchanged for MilliQ water and the gels incubated for 30 min, with two more exchanges of water each lasting 30 min. The gels were stored at 4°C over night. The stained gel pieces were mounted onto glass bottom imaging dishes (35 mm Glass Bottom µ-Dish, ibidi, 81158) by overlaying with 2% low melting-point agarose (Promega, V3841) and either imaged directly or stored at 4°C until imaging.

**Table 2:** Lists of primary antibodies used in this study.

| Primary antibodies | Antibody type | Antibody source | Catalogue number | Antibody dilution |
| --- | --- | --- | --- | --- |
| ALIX [EPR23653-32] | Rabbit monoclonal | Abcam | ab275377 | 1:200 |
| ALIX | Mouse polyclonal | Abcam | ab88743 | 1:200 |
| Beta-tubulin (2-28-33) | Mouse monoclonal | Sigma Aldrich | T5293 | 1:400 |
| Alpha-tubulin (B-5-1-2) | Mouse monoclonal | Sigma Aldrich | T5168 | 1:400 |
| Alpha-tubulin [YL1/2] | Rat monoclonal (IgG2a) | Abcam | ab6160 | 1:400 |
| Anillin | rabbit | Gift from Christine Field |  | 1:200 - 1:250 |
| Beta-actin | Mouse monoclonal (IgG2b) | Proteintech | 66009-1-Ig | 1:400 |
| CDC42EP4 (BORG4) | Rabbit polyclonal | Prestige Antibodies/ Sigma Aldrich | HPA023335 | 1:200 |
| CHMP4B | Rabbit polyclonal | Abcam | ab105767 | 1:100 |
| CRIK/Citron Kinase | Mouse monoclonal (IgG1) | BD Biosciences | 611377 | 1:400 |
| GFP | Goat polyclonal | Abcam | ab6673 | 1:200 |
| hIST1 | Rabbit polyclonal | Proteintech | 51002-1-AP | 1:50 |
| MKLP1 [N1N2] | Rabbit polyclonal | GeneTex | GTX120875 | 1:250 |
| Non-muscle Myosin Heavy Chain II-A | Rabbit polyclonal | Convance<br>(now Biolegend) | PRB-440P<br>(909801) | 1:200 |
| Non-muscle Myosin Heavy Chain II-B | Rabbit polyclonal | Convance<br>(now Biolegend) | PRB-445P<br>(909901) | 1:300 |
| PRC1 (16F2) | Mouse monoclonal (IgG2b) | Thermo Fisher | MA1-846 | 1:500 |
| RACGAP1 | Rabbit polyclonal | Proteintech | 13739-1-AP | 1:500 |
| Septin11 | rabbit | Kind gift from Bill Trimble |  | 1:200 |
| hSeptin7 (C) | Rabbit polyclonal | IBL | JP18991 | 1:200 |
| Septin9 | Rabbit polyclonal | Novus Biologicals | NBP2-13294 | 1:200 |
| TSG101 | Rabbit polyclonal | Proteintech | 28283-1-AP | 1:200 |
| TSG101 [4A10] | Mouse monoclonal (IgG1) | Abcam | ab83 | 1:200 |

**Table 3:**
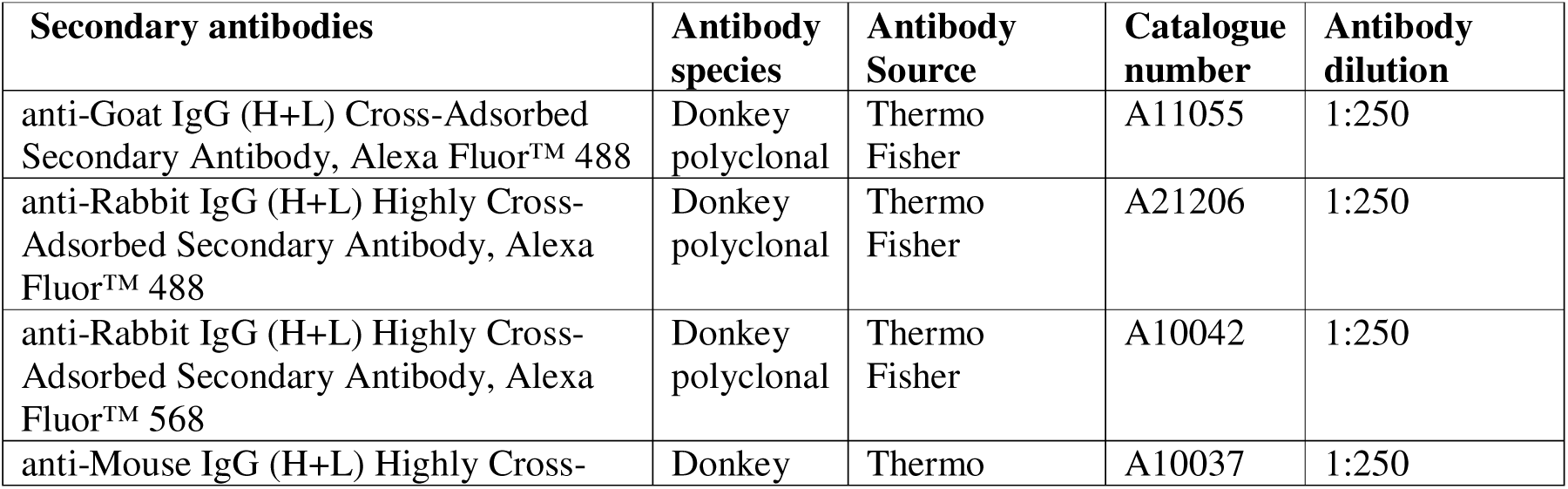
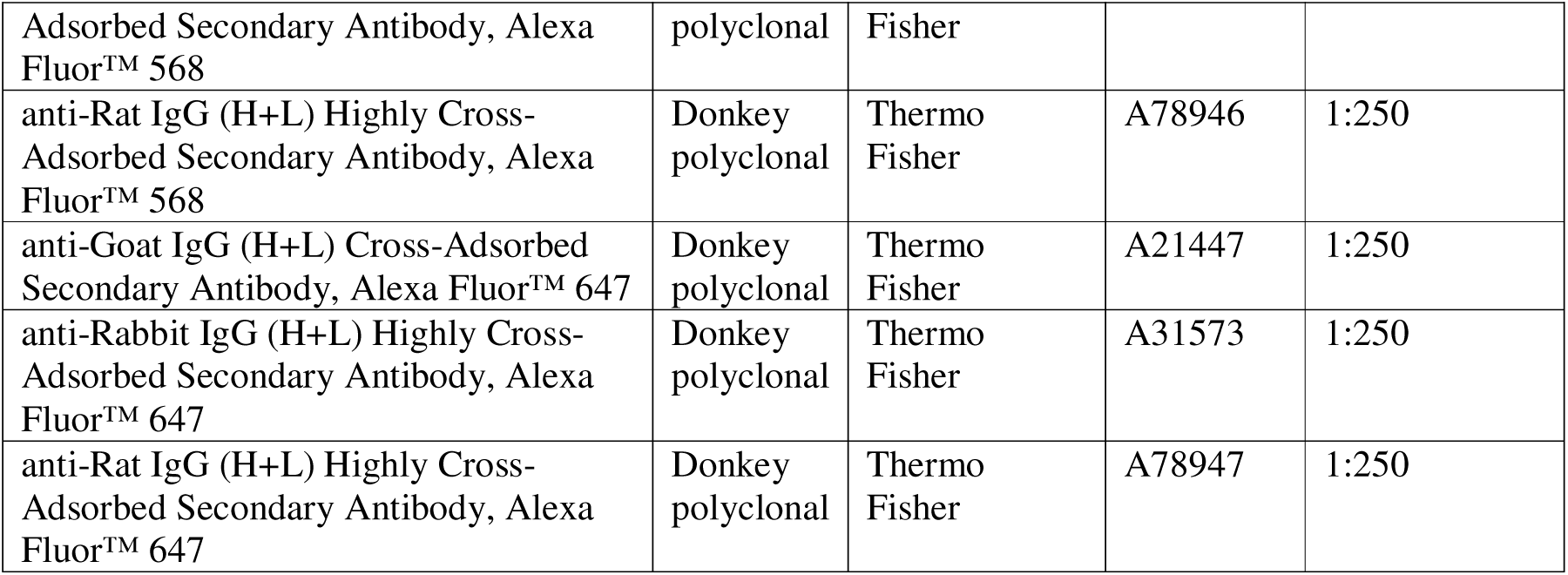
Lists of secondary antibodies used in this study.

**Table 4:** List of stains used in this study.

| Stains | Stain Source | Catalogue number | Stain dilution/concntration |
| --- | --- | --- | --- |
| DAPI (4',6-Diamidin-2-phenylindol, Dihydrochlorid) (5 mg/ml) | Thermo Fisher | D1306 | 1:500 |
| Hoechst 33342, trihydrochloride, trihydrate (10 mg/ml) | Thermo Fisher | H3570 | 1:500 |
| CellMask Orange Plasma Membrane Stain (5 mg/ml) | Thermo Fisher | C10045 | 1:1000 |
| SiR-tubulin | Spirochrome | SC002 | 100 nM |

### Pre-and post-expansion comparison

Synchronized and fixed cells were processed with the U-ExM anchoring solution as above, followed by an immunofluorescence staining. The anchored samples were washed thrice for 10 min with PBS. The cells were permeabilized with 0.3% v/v Triton-X100 in PHEM buffer for 5 min, and briefly washed with PHEM buffer. Next, they were blocked with 4% v/v horse serum (Gibco, 160555130), 1% v/v BSA in PHEM buffer for 1 h. Primary antibody against alpha-tubulin (Sigma Aldrich, T5168) was diluted 1:400 in 1% v/v BSA/PHEM and incubated with the coverslips for 2 h. The coverslips were washed thrice with PHEM buffer for 10 min, followed by incubation with secondary donkey anti-mouse IgG antibody coupled to Alexa Fluor 568 (Thermo Fisher, A10037), diluted 1:400 in 1% v/v BSA/PHEM for 1 h.

The coverslips were washed thrice with PHEM buffer for 10 min. The stained coverslips were imaged and directly transformed into gels as described above. The expanded gels were stained with primary antibodies against alpha-tubulin (Abcam, ab6160) and GFP (Abcam, ab6673) and secondary antibodies donkey anti-goat IgG Alexa Fluor 488 (Thermo Fisher, A11055) and donkey anti-rat IgG Alexa Fluor 568 (Thermo Fisher, A78946), all diluted in 2% BSA/PBS (see *Staining of U-ExM gels* for a more detailed protocol). The gels were imaged with a z-spacing of 0.5 µm. To compare the same cell before and after expansion, maximum intensity projections of the z-stacks were compared with the bf_template_match script (as used in Reference ^68^, https://github.com/sommerc/expansion_factor_bftm). The initial expansion factor was set to 3.8-4.5 with a step size of 0.1; the rotation angle was set from 1-360 degrees, with a step size of 1 degree. The pixel size of the pre-and post-expansion image corresponded to 90 nm. All other values were kept as default. The best matching pairs of pre-and post-expansion images were further analyzed for distortion as described^68^.

### Expansion factor calculation

All gels were measured with a caliper after expansion and the expansion factor (EF) was calculated as the ratio of the expanded gel and the original coverslip size. Of all gels used for this study, the average EF was calculated and applied to convert pixel sizes.

### Lattice light-sheet image acquisition

Time-lapse movies of dividing cells were acquired on a Lattice Light-Sheet system (LLS7, Zeiss), equipped with 488 nm and 640 nm lasers and a sCMOS camera (pco.edge 4.2 CLHS). In the LLS7 the sample is scanned through a stable (oblique) light sheet and the rate at which volume scans are acquired vary depending on the scan distance (FOV), the step-size of the scan and the exposure time. The effective interval at which a single volume scan was acquired varied therefore between 5 min and 5 min 35 sec. We used the light-sheet Sinc3 30×1000 with a fixed length of 30 µm and thickness of 1 µm, a variable scanning distance between 650 and 1000 µm and a step-size of 0.2 µm. The volumetric movies were acquired at 37°C, 5% CO_2_ in humidified atmosphere for 10-12 h. In the LLS7, the sample is illuminated at an angle of 30° and the emission is detected at an angle of 60° relative to the coverslip.

Therefore, the raw images were processed using the features ‘cover glass transformation’ and ‘deskew’ of ZEN Blue Software (Zeiss, Version 3.5), resulting in distortion-corrected ‘deskewed’ volumes with the conventional orientation in x/y/z in relation to the coverslip.

### Imaging of fixed samples and U-ExM gels

All images of expanded samples and fixed cells on coverslips were acquired with an inverted Olympus IX71 microscope equipped with a CSU-X1 spinning disk unit (Yokogawa) and a ORCA Flash 4.0LT sCMOS camera (Hamamatsu). A 60×/1.42 NA oil Olympus objective was used together with a 405 nm (100 mW; VORTRAN), a 491 nm (100 mW; Cobolt) a 561 nm (100 mW; Cobolt), and a 639 nm (VORTRAN, 160 mW) laser. Lasers were controlled by an iLas laser illumination system (Gataca Systems) and operated with the software MetaMorph (Molecular Devices). Unless otherwise indicated, z-stacks with a spacing of 1 µm were acquired of the gels. For fixed unexpanded samples, the z-interval was 0.5 µm.

## DATA ANALYSIS

Analysis of cytokinetic stages in lattice light-sheet movies.

A total of 206 cell-division events were extracted and inspected from 7 lattice light-sheet movies. 129 movies, which depicted the full process from cleavage furrow formation until abscission as judged by microtubule severing, were analyzed further. Five signature events that occurred in all of the movies were identified: 1) Septin assembly at the cleavage furrow, 2) constriction of the daughter cells, 3) spreading of the daughter cells, 4) splitting of the septin rings and 5) abscission. To measure the time until the onset of these events, the assembly of septins at the cleavage furrow was defined as t=0. The diameter of the microtubule bundle was measured with the line scan tool in FIJI^75^. A line with a width of 25 pixel was drawn orthogonally to the ICB and the resulting intensity profile was approached with a Gaussian fit. The full width at half maximum (FWHM) of the Gaussian fit was extracted. The last frame before abscission (as defined by microtubule severing) was measured and added as ‘before abscission’ to the dataset.

### Fourier Ring Correlation

To estimate the resolution achievable with the U-ExM protocol, samples were stained against alpha-tubulin as described above. Two images of the same cell were acquired and analyzed with the Fourier ring correlation tool by Alex Herbert (https://github.com/BIOP/ijp-frc) based on Reference ^76^. The three sigma method was used to determine the threshold for high frequency noise. A total of 20 images were analyzed.

### Measurement of line profiles

Line profiles were measured on maximum intensity projections of individual U-ExM images, normalized and averaged. Line profiles were measured as indicated in the main text, dependent on the area that was to be inspected: 1) To assess target distribution along the ICB axis, a 25 pixel-wide line was drawn in FIJI along the cytokinetic axis, with the centroid of the line placed in the center of the midbody. Orientation of this line was normalized to account for asymmetric abscission. 2) To assess the target distribution throughout the midbody, a 25 pixel-wide line was drawn across the midbody, with its centroid on the edge of the microtubule bundle. The line was oriented so that the outside of the midbody always aligned with the same end of the line. 3) To assess the target distribution on the ICB arm, a 25 pixel-wide line was drawn across the ICB arm, with its centroid on the edge of the microtubule bundle. The line was oriented so that the outside of the bridge always aligned with the same end of the line.

### Image Processing and Feature Extraction with CellProfiler

To analyze the volumetric multichannel U-ExM dataset of 20 targets in pseudotime, we developed a customized feature extraction pipeline using CellProfiler 4^77^. Prior to analysis with CellProfiler, the z-stacks were projected using maximum intensity projection, and the z-projected images were pre-processed with a custom ImageJ macro that split multichannel stacks into separate grayscale image sequences per channel. Only the microtubule and Septin2-GFP channels were used as inputs for the CellProfiler pipeline. Among these, the microtubule channel served as the primary basis for segmentation, then used for both morphological and intensity-based feature extraction, while the Septin2 channel provided additional intensity-based context. The input images were loaded, structured and organized using the *Images, Metadata, NamesAndTypes,* and *Groups* modules. Channel-specific normalization and contrast enhancement were done with the *RescaleIntensity* and *ImageMath* modules. After contrast enhancement and rescaling, the ICB structures were segmented in the microtubule channel using the *IdentifyPrimaryObjects* module in CellProfiler. The module was configured to detect bridge-like structures with a typical diameter range of 50 to 400 pixels, reflecting the empirically observed size range of the structures of interest in the dataset. To ensure precision, objects that were very close to or touching the image borders, as well as those falling outside the expected diameter range were discarded. An adaptive thresholding strategy was applied, using a two-class Otsu segmentation method to distinguish foreground from background within each image. An adaptive window size of 50 pixels was used to locally compute the threshold, improving segmentation in images with uneven illumination or complex background. No additional declumping method was applied at this stage, as bridges were typically well-separated or followed up by post-filtering modules for refinement. The *FilterObjects* was used for morphological refinement ensuring robust segmentation and minimizing false positives by eliminating small artifacts and enforcing size/shape thresholds. The segmentation logic was fine-tuned across datasets using an iterative combination of manual inspection and grid-search-based parameter adjustment to achieve a balance between sensitivity and precision. While we aimed to apply consistent parameter settings across all targets, occasional target-specific minor adjustments and manual corrections were necessary due to inter-target variability. For feature extraction, we combined object-level measurements and image-level measurements. Object-level measurements included *MeasureObjectSizeShape, MeasureObjectIntensity,* and *MeasureTexture* (computed over multiple scales). For image-level measurements *MeasureImageIntensity* was used to capture global signal statistics. *ExportToSpreadsheet* was used to generate both object-level and image-level feature tables. *SaveImages* exported overlaid and labeled masks for manual curation. To prepare features for downstream analysis, we implemented several Python scripts using Python 3.10 with *pandas*^78^*, numpy*^79^ and *openpyxl.* The largest object per image was retained (using *AreaShape_Area* feature) to focus on the dominant bridge for size and morphology measurements. We also recorded the total object count per frame. Object-level features were grouped by image. This generated one representative row per frame. Finally. object-level and image-level features were merged.

### Feature selection

For pseudotime alignment of stages, we combined the image features extracted via CellProfiler that described the morphology of the masked ICB with a set of manually extracted features that refined our description of the microtubule bundle and Septin2-GFP distribution along the ICB.

The CellProfiler-based features were descriptors of the 2D shape of the developing ICB, which is represented by a mask in the tubulin channel: 1) *areashape_equivalentdiameter;* the diameter of a circle with the same area as the masked ICB, a descriptor of the area size of the ICB which decreases as cytokinesis progresses and the central spindle becomes more and more compact, 2) *areashape_maximumradius;* the maximum distance of any pixel in the masked ICB to the closest pixel outside of the object, a descriptor of the shape of the ICB and its eccentricity, which changes especially in early cytokinetic steps, 3) *areashape_meanradius*; the mean distance of any pixel in the masked ICB to the closest pixel outside of the object, which decreases with shrinking of overall tubulin-positive area during cytokinetic progression, 4) *areashape_eccentricity;* the eccentricity of the ellipse that has the same second-moments as the masked ICB. As the ICB grows thinner and over time, the ellipse describing it becomes more eccentric, 5) *areashape_minoraxislength;* the length of the minor axis of the ellipse that has the same normalized second central moments as the region, which reflects the thinning of the ICB, 6) *intensity_integratedintensity_input_septin;* the intensity of the septin staining in the masked area, which changes as septins move from the ingressed furrow to their ring and manchette organization and disappear from the ICB as abscission approaches and 7) *intensity_integratedintensity_input_microtubule;* the intensity of the tubulin staining in the masked area, which decreases as the microtubule bundle becomes thinner and microtubules are disassembled as abscission approaches. A ranked list of these features and their contribution to PC1 and PC2 can be found in Table 1.

The manually measured features included 1) the diameter of the midbody, 2) the diameter of the ICB arm, which is defined as the narrowest position on the microtubule bundle, 3) the ratio of the these two diameters, 4) the distance between the Septin2-GFP structures along the major axis of the ICB and, 5) a Boolean feature to describe whether the microtubule bundle in the ICB was cut or not. The diameter of the midbody and ICB were measured in FIJI, by drawing an orthogonal 25 pixel-wide line across the tubulin-based structures and approaching the resulting curve with a Gaussian fit. The FWHM of the Gaussian curve was stored and corrected for the expansion factor. The diameters of the midbody, the ICB arm and their derivatives were used to determine the degree of thinning of the microtubule ICB. The relative distribution of Septin2-GFP along the bridge helped determine the stage of cytokinesis, as we found that septin moved gradually from the center of the midbody to the ICB-cell junctions throughout the process.

### Principle component analysis of aggregated features for pseudotime alignment

Principal Component Analysis (PCA) was applied to selected features for dimensionality reduction. We normalized all features via z-score prior to feeding them to the PCA algorithm. The first two principal components were selected to represent the images. The manually assigned stages were color-coded for comparison. We then constructed a 4^th^ degree polynomial fit of the 2D points in PCA space and collapsed all points to this fit line. Moving along this line created a continuous representation of pseudotime space, which allowed a chronological sorting of the pre-assigned stages. Because the fit line is continuous, it is possible to divide into an arbitrary number of equally sized pseudotemporal bins, which were used to generate a 34 frame-long movie of the cytokinesis. We used a step size of 49 with an overlap of 49, resulting in a bin size of 98 images. All dimensionality reduction was performed with custom written Python scripts using *scikit-learn*^80^, and visualizations were generated with Python libraries *matplotlib*^81^ and *seaborn*^82^.

### Cytokinetic pseudotime image reconstruction

Each raw image was a volumetric z-stack with 4 color channels. These included a channel for imaging of microtubules, a channel for Septin2-GFP, a channel for the nucleus (DAPI), and an additional target protein. Images were averaged per cytokinetic stage and per target using the full z-stacks, radial averages, or mean intensity projections.

To align the ICBs, the bridge axis was defined in FIJI. Maximum intensity projections were created and a line was drawn along the ICB axis. The centroid was placed on the midzone of the bridge. In case of asymmetrically abscised bridges, the orientation of the line was kept similar among all the images (i.e. the positive endpoint was placed on the cell-connected arm of the bridge, while the negative endpoint was located towards the abscission site). The line was defined by its centroid, length and angle. The original image volumes were aligned in the lateral plane using these line parameters. Each volume was shifted so that the center of the line corresponded to the center of the lateral field of view in the stack.

Each volume was then cropped in the lateral plane to a fixed size, given by √2\**length*, where *length* (usually, 500 pixels) was the target size of the region to crop around the midbody. The factor √2 ensured the volume could be cropped to a size *length* without losing any data, even after rotation by up to 45 degrees.

Each volume was then rotated according to the angle of the line drawn along the major axis of the ICB, with respect to the vertical axis of the lateral plane, so that the bridge lay straight with respect to the second to last axis of the image. Rotation was done with nearest neighbor interpolation to ensure fast computation.

Each volume was then normalized to its minimum and maximum values, so all counts were in the range [0,1].

Following this, if the image was to be used for z-stacking or radial projection, a copy of this volume was resliced along an axial (xz) plane, to get an orthogonal view of the midbody. This was then cropped to the central 25 x-pixels of the major axis of the bridge. A Gaussian profile was fit along the axial direction (z) to the mean projection along the orthogonal lateral direction (y) of the 25 x-pixels. The mean value of the fit Gaussian was taken to be the vertical center, termed the central z-coordinate, of the midbody.

The normalized volume was then cropped in the lateral plane to a size *length*. If two distinct septin rings were clearly found in all images in a specific cytokinetic stage, each image was rescaled so that the distance between the rings was the same. To do so, we measured the intensity of the septin channel along the ICB axis and looked for local maxima. The two highest intensity values were defined as the septin ring location. This helped ensure better overlap of multiple images.

If the image was to be used for z-stacking or radial projection, it was rescaled in the axial (z) direction according to the ratio of the z-sampling (usually, 1 µm) over the pixel size of the camera (usually, 90 nm), which was ∼11.1x. This improved the visual interpretability of the mean projections. Upsampling was performed using a nearest neighbor interpolation to ensure fast computation.

Following this, multiple images of targets were summed to create an average image. For z-stack averages, images were added to an initially empty volume such that their fit central z-coordinate lay at the axial center of the volume (length of the z-dimension of the empty volume, divided by 2). This resulted in one 3D stack per target. For radial averages, a cylindrical coordinate space was constructed so that its center lay along the major axis of the microtubule bridge. Concentric shells of this sphere in thicknesses specified by *bin_size* (usually 1 pixel) were summed, normalized by their voxel count, and placed as single rows in a resulting 2D image showing the projection of the signal in the original volume at this radius. This resulted in one 2D image per target. For mean projections, the images were summed along their axial direction and then added together to create one 2D image per target.

Image normalization, translation and rotation was performed in Python using the *numpy*^79^ and *scipy*^83^ libraries. Code availability.

The original software for pseudotime binning and image reconstruction is provided as open source with a BSD-3 license, available online at https://github.com/zacsimile/cytokinesis-pseudotime-analysis

The original software for image feature extraction is provided as open source with a MIT license, available online at https://github.com/AG-Ewers/cytokinesis-feature-analysis

### Statistics

A comprehensive list of the number of images that were used to generate the averaged representations shown in Figures 3, 4, 5, 6, 7 and 8 can be found in Table 5.

**Table 5:**
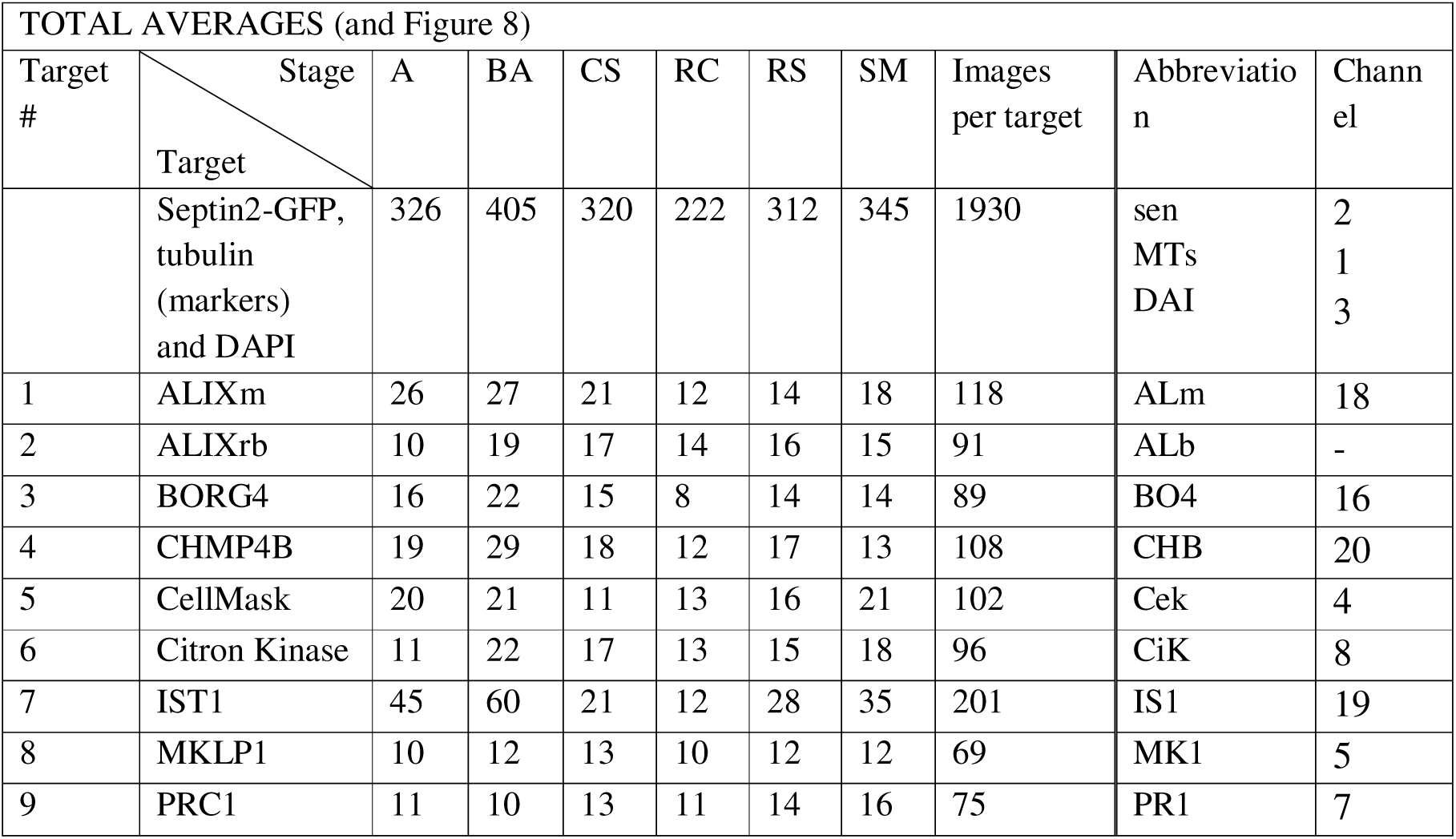
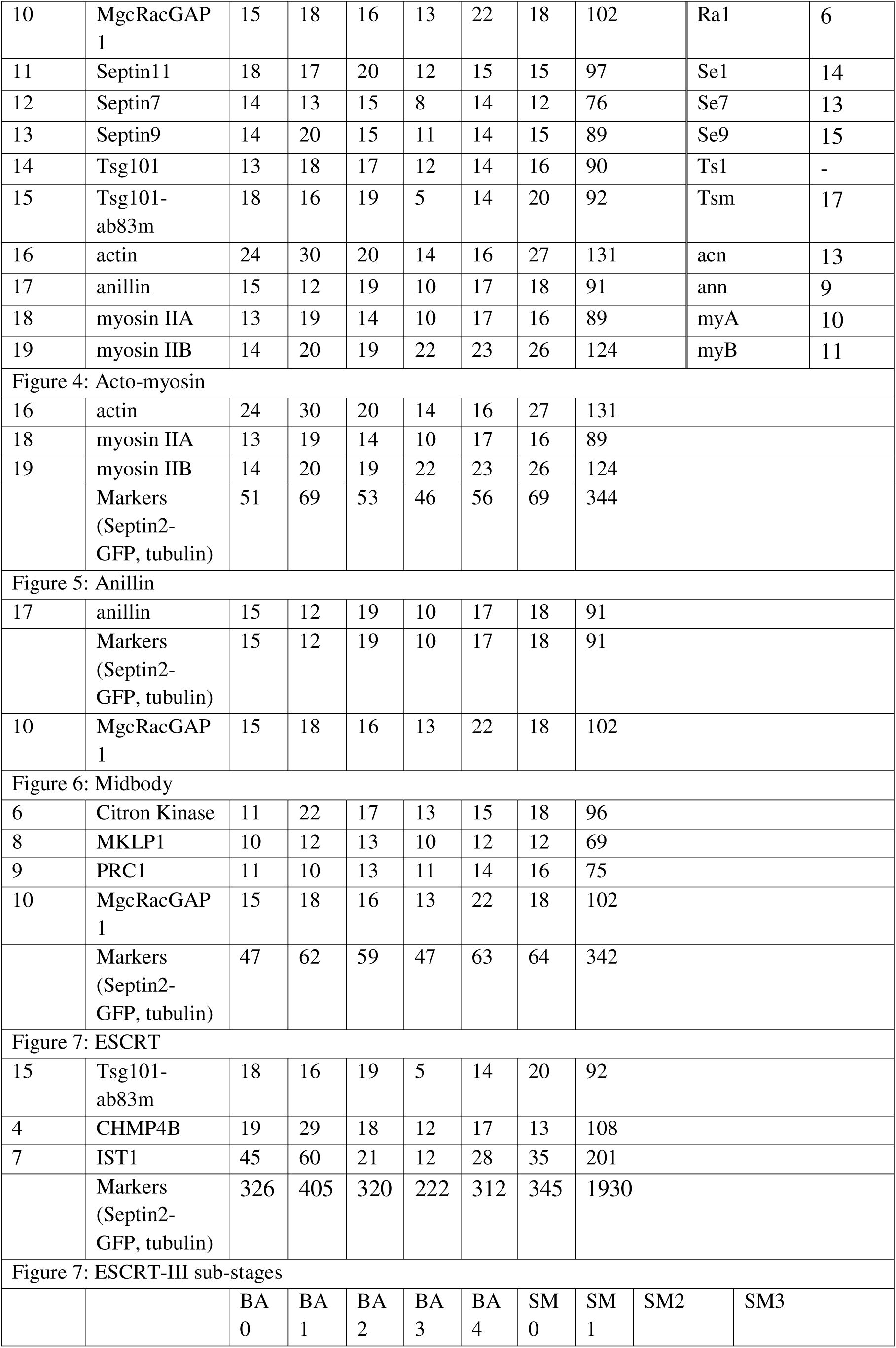
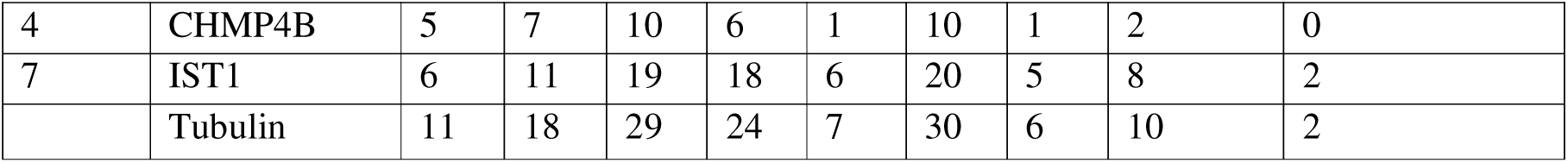
Number of images used for pseudotime image reconstruction of cytokinesis.

| TOTAL AVERAGES (and Figure 8) |  |  |  |  |  |  |  |  |  |  |
| --- | --- | --- | --- | --- | --- | --- | --- | --- | --- | --- |
| Target # | Stage<br>Target | A | BA | CS | RC | RS | SM | Images per target | Abbreviation | Channel |
|  | Septin2-GFP, tubulin (markers) and DAPI | 326 | 405 | 320 | 222 | 312 | 345 | 1930 | sen<br>MTs<br>DAI | 2<br>1<br>3 |
| 1 | ALIXm | 26 | 27 | 21 | 12 | 14 | 18 | 118 | ALm | 18 |
| 2 | ALIXrb | 10 | 19 | 17 | 14 | 16 | 15 | 91 | ALb | - |
| 3 | BORG4 | 16 | 22 | 15 | 8 | 14 | 14 | 89 | BO4 | 16 |
| 4 | CHMP4B | 19 | 29 | 18 | 12 | 17 | 13 | 108 | CHB | 20 |
| 5 | CellMask | 20 | 21 | 11 | 13 | 16 | 21 | 102 | Cek | 4 |
| 6 | Citron Kinase | 11 | 22 | 17 | 13 | 15 | 18 | 96 | CiK | 8 |
| 7 | IST1 | 45 | 60 | 21 | 12 | 28 | 35 | 201 | IS1 | 19 |
| 8 | MKLP1 | 10 | 12 | 13 | 10 | 12 | 12 | 69 | MK1 | 5 |
| 9 | PRC1 | 11 | 10 | 13 | 11 | 14 | 16 | 75 | PR1 | 7 |
| 10 | MgcRacGAP 1 | 15 | 18 | 16 | 13 | 22 | 18 | 102 | Ra1 | 6 |
| 11 | Septin11 | 18 | 17 | 20 | 12 | 15 | 15 | 97 | Se1 | 14 |
| 12 | Septin7 | 14 | 13 | 15 | 8 | 14 | 12 | 76 | Se7 | 13 |
| 13 | Septin9 | 14 | 20 | 15 | 11 | 14 | 15 | 89 | Se9 | 15 |
| 14 | Tsg101 | 13 | 18 | 17 | 12 | 14 | 16 | 90 | Ts1 | - |
| 15 | Tsg101-ab83m | 18 | 16 | 19 | 5 | 14 | 20 | 92 | Tsm | 17 |
| 16 | actin | 24 | 30 | 20 | 14 | 16 | 27 | 131 | acn | 13 |
| 17 | anillin | 15 | 12 | 19 | 10 | 17 | 18 | 91 | ann | 9 |
| 18 | myosin IIA | 13 | 19 | 14 | 10 | 17 | 16 | 89 | myA | 10 |
| 19 | myosin IIB | 14 | 20 | 19 | 22 | 23 | 26 | 124 | myB | 11 |

|  |  |  |  |  |  |  |  |  |
| --- | --- | --- | --- | --- | --- | --- | --- | --- |
| 16 | actin | 24 | 30 | 20 | 14 | 16 | 27 | 131 |
| 18 | myosin IIA | 13 | 19 | 14 | 10 | 17 | 16 | 89 |
| 19 | myosin IIB | 14 | 20 | 19 | 22 | 23 | 26 | 124 |
|  | Markers (Septin2-GFP, tubulin) | 51 | 69 | 53 | 46 | 56 | 69 | 344 |

|  |  |  |  |  |  |  |  |  |
| --- | --- | --- | --- | --- | --- | --- | --- | --- |
| 17 | anillin | 15 | 12 | 19 | 10 | 17 | 18 | 91 |
|  | Markers (Septin2-GFP, tubulin) | 15 | 12 | 19 | 10 | 17 | 18 | 91 |
| 10 | MgcRacGAP 1 | 15 | 18 | 16 | 13 | 22 | 18 | 102 |

|  |  |  |  |  |  |  |  |  |
| --- | --- | --- | --- | --- | --- | --- | --- | --- |
| 6 | Citron Kinase | 11 | 22 | 17 | 13 | 15 | 18 | 96 |
| 8 | MKLP1 | 10 | 12 | 13 | 10 | 12 | 12 | 69 |
| 9 | PRC1 | 11 | 10 | 13 | 11 | 14 | 16 | 75 |
| 10 | MgcRacGAP 1 | 15 | 18 | 16 | 13 | 22 | 18 | 102 |
|  | Markers (Septin2-GFP, tubulin) | 47 | 62 | 59 | 47 | 63 | 64 | 342 |

|  |  |  |  |  |  |  |  |  |
| --- | --- | --- | --- | --- | --- | --- | --- | --- |
| 15 | Tsg101-ab83m | 18 | 16 | 19 | 5 | 14 | 20 | 92 |
| 4 | CHMP4B | 19 | 29 | 18 | 12 | 17 | 13 | 108 |
| 7 | IST1 | 45 | 60 | 21 | 12 | 28 | 35 | 201 |
|  | Markers (Septin2-GFP, tubulin) | 326 | 405 | 320 | 222 | 312 | 345 | 1930 |
|  |  | BA 0 | BA 1 | BA 2 | BA 3 | BA 4 | SM 0 | SM 1 | SM2 | SM3 |
| 4 | CHMP4B | 5 | 7 | 10 | 6 | 1 | 10 | 1 | 2 | 0 |
| 7 | IST1 | 6 | 11 | 19 | 18 | 6 | 20 | 5 | 8 | 2 |
|  | Tubulin | 11 | 18 | 29 | 24 | 7 | 30 | 6 | 10 | 2 |

Statistical tests and data visualization were carried out in OriginPro 2025 (OriginLab Coorporation, USA).

## Acknowledgements

We thank all the Ewers laboratory for helpful discussions. We thank Olivia Heese for help in the early stage of data analysis development, Andrea Senge for help with cell synchronization. Lattice-light sheet data were acquired in the AMBIO microscopy facility at Charité Universitätsmedizin Berlin. We thank Dr. Stefan Donat for help with the microscopy at AMBIO. We would like to acknowledge the Core Facility BioSupraMol supported by the DFG. The work was funded by Freie Universität Berlin and Deutsche Forschungsgemeinschaft (DFG) through SFB958 and project no. 278001972 - TRR 186.

## Author contributions

HE and NH conceived the project. HE and NH designed experiments. NH and KB performed experiments. NH, KB, ZM, AZ, DS and JR performed image analysis. JS setup the LLS microscopy. NH and HE wrote the paper with input from all authors.

---

**Supplementary Figure S1:**
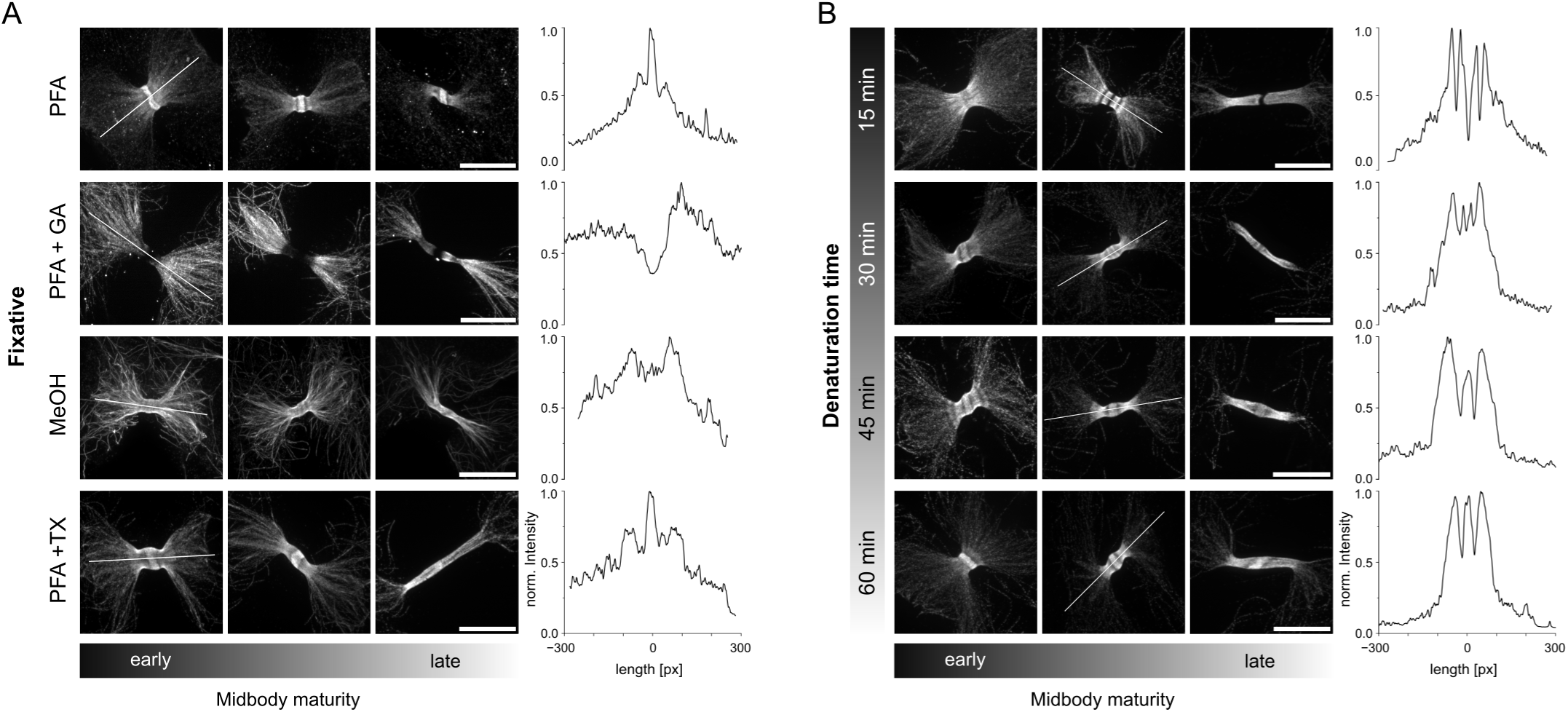
Optimization of U-ExM protocol to accommodate full midbody staining. A) Comparison of the effect of fixatives (PFA: 4 % paraformaldehyde, PFA + GA: 4 % paraformaldehyde with 0.05 % glutaraldehyde, MeOH: methanol, PFA + TX: 4 % paraformaldehyde with 0.1% Triton X-100,) on midbody accessibility for antibodies. Immunofluorescence staining against alpha-Tubulin. Three different stages of midbody development are shown. Line profiles on the right are taken from the early developing midbodies. B) Correlation of midbody accessibility for antibodies with the denaturation of the hydrogel. Hydrogels were denatured for either 15, 30, 45 or 60 minutes and immunostained against alpha-Tubulin. Three different stages of midbody development are shown. Line profiles on the right are taken from the intermediate developing midbodies. All images are maximum intensity projections. Scale bar is 5 µm.

**Supplementary Figure S2:**
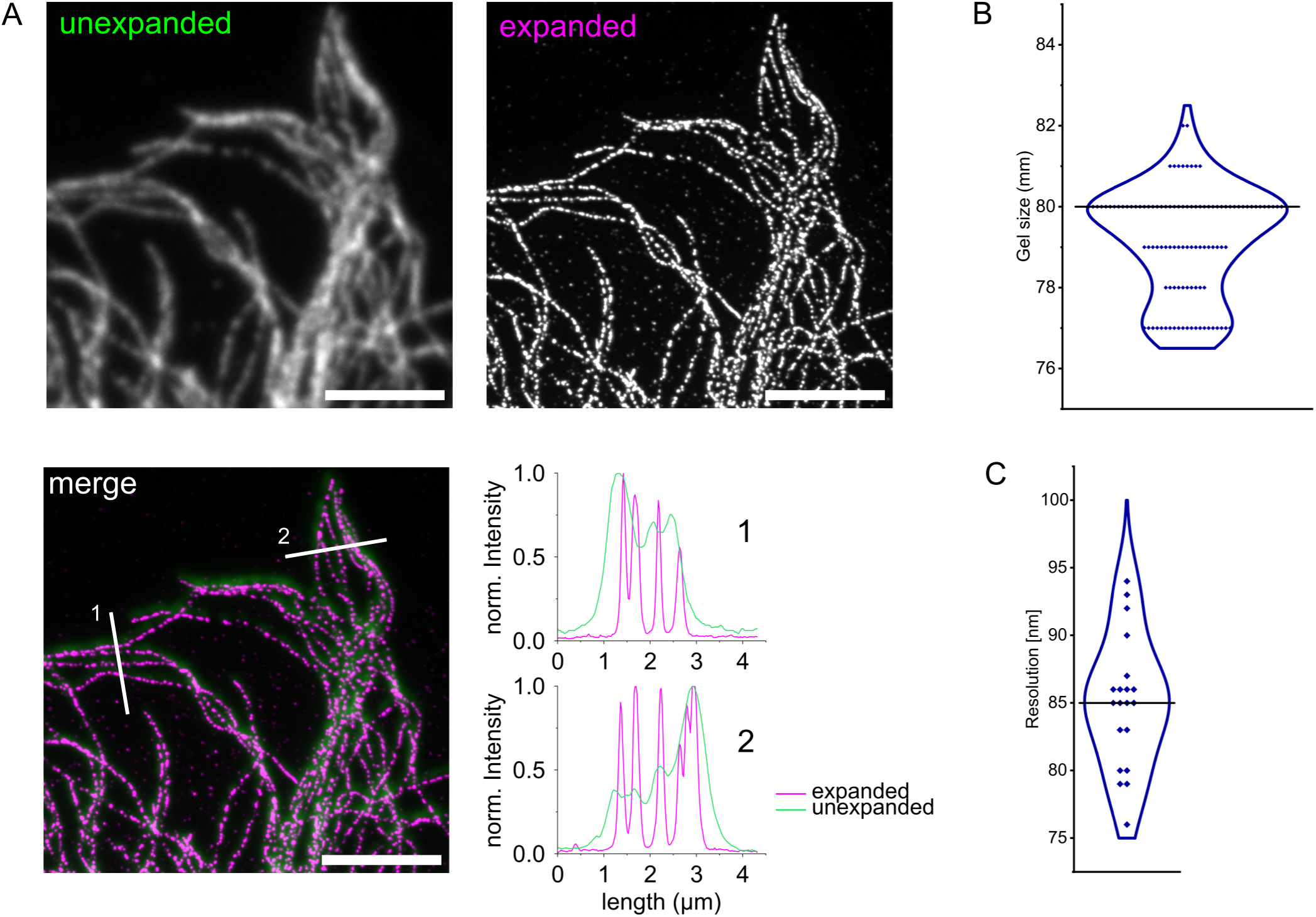
Control of resolution of U-ExM. A) Immunofluorescence images of cell stained against alpha-Tubulin before (green) and after expansion (magenta). Line profiles across two different areas (1 and 2) show the increased resolution in the expanded image. Scale bars are 5 µm and 21.5 µm in the unexpanded and the expanded sample, respectively. All images are maximum intensity projections. B) Diameter of the expanded gels that were used for this study. An average gel size of 80 (marked by black solid line) results in an expansion factor of 4.44. A total of n=109 gels was used. C) Resolution of expanded images based on Fourier ring correlation analysis using the three sigma threshold of n=20 samples with immunofluorescence staining against alpha-Tubulin. The mean resolution is marked (black solid line).

**Supplementary Figure S3:**
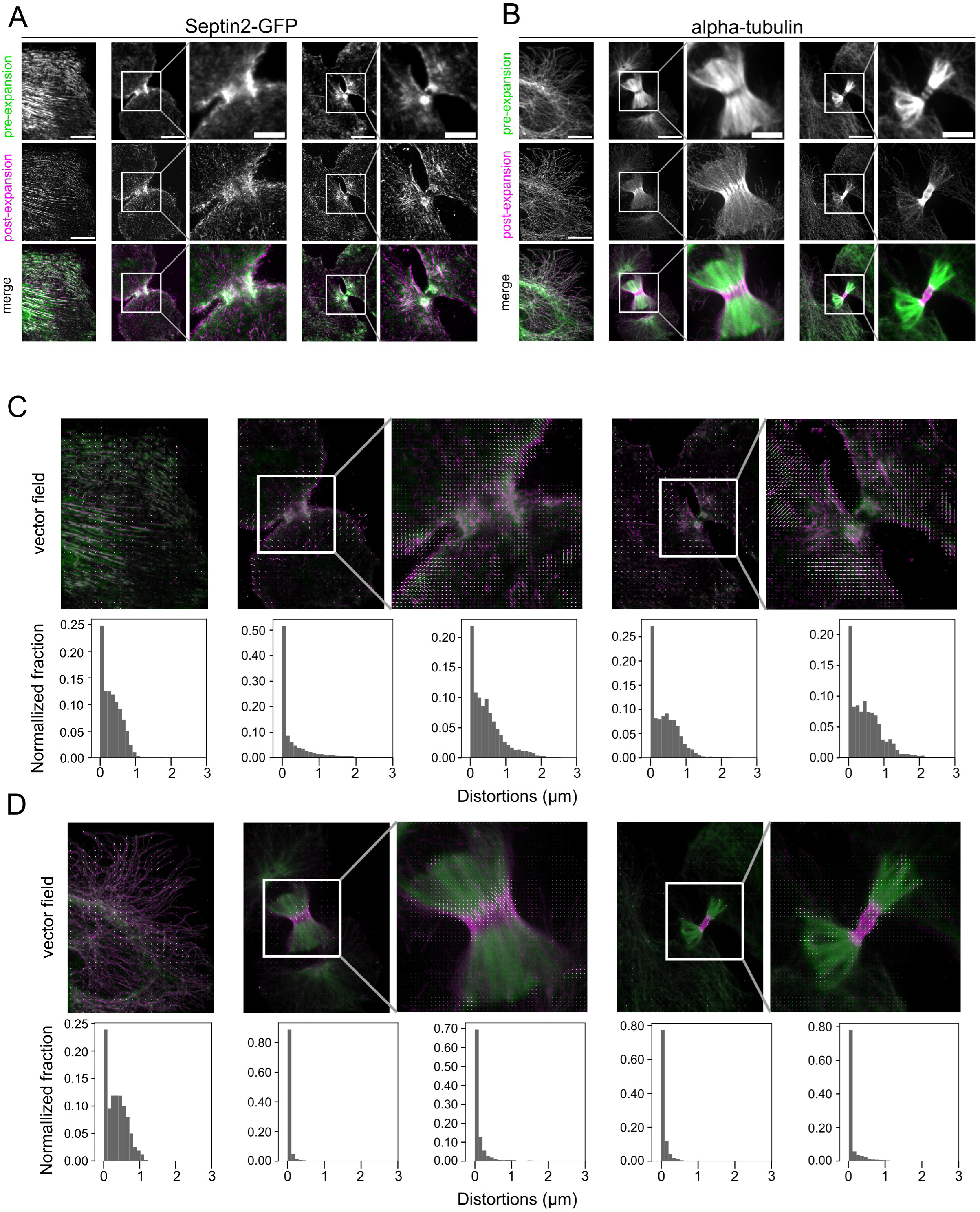
Quality control of distortion in U-ExM. A) Comparison of the pre-expansion (green) and post-expansion (magenta) images of three different cells stained for Septin2-GFP. One interphase cell, one cleavage furrow and one ICB are shown with zooms into the spindle and midbody area (marked by white squares in overview images). Merges show the similarity transformed pre-and post-expansion images. B) Same as in A) but the alpha-Tubulin staining is shown. C) Distortion analysis for Septin2-GFP. Vector fields illustrate the direction and amount of distortion between the transformed pre-and post-expansion images. The histograms below show the normalized fraction of the distortion vectors sorted by length. D) Same as in D) but the alpha-Tubulin staining is shown. All images are maximum intensity projections. Scale bars in A) and B) are corrected for the expansion factor and represent 10 µm in overview images and 5 µm in the zoomed in areas.

**Supplementary Figure S4:**
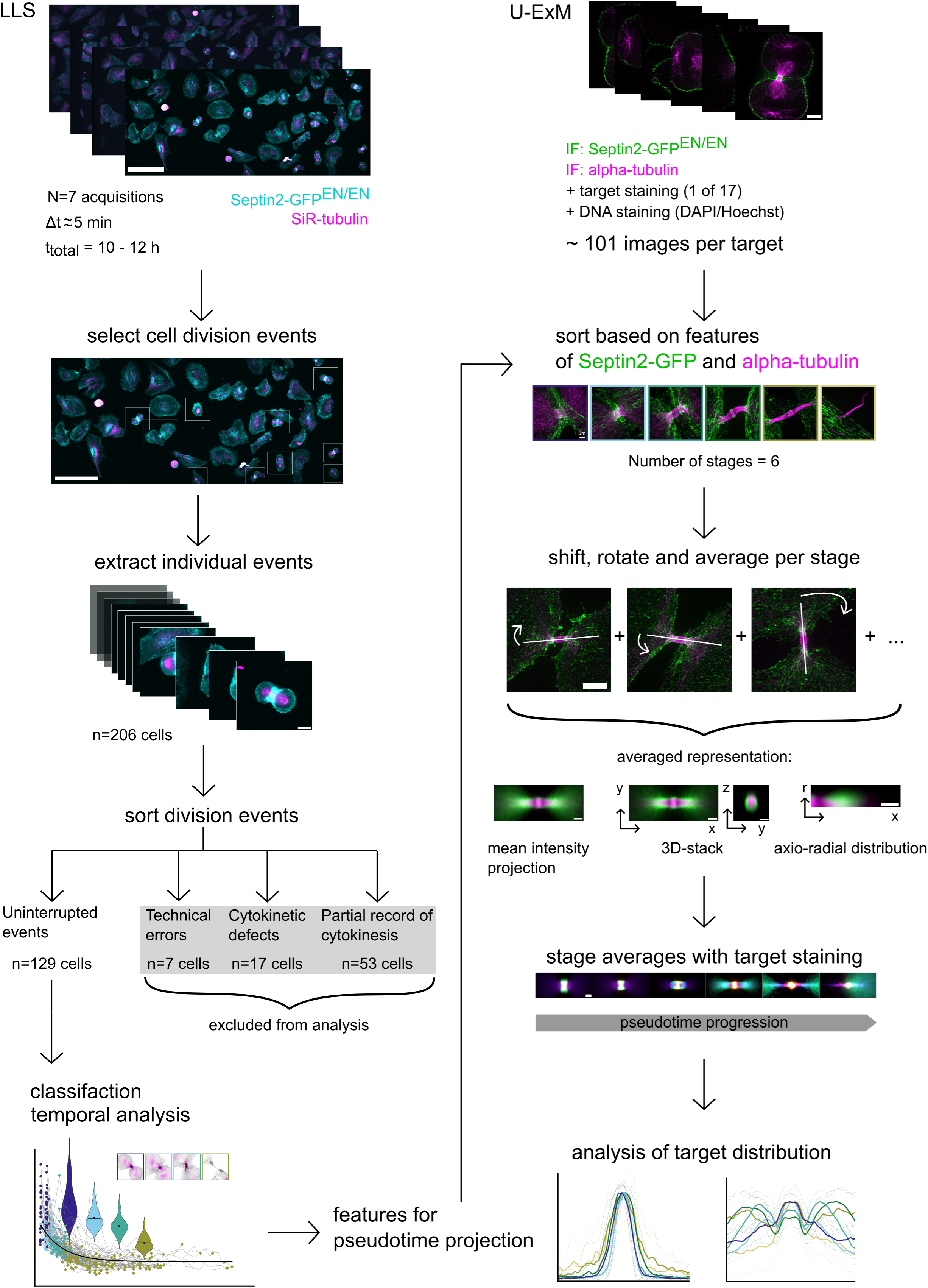
Experimental strategy to correlate lattice-light sheet and U-ExM imaging. Left: Experimental workflow for the lattice light sheet data acquisition. N=7 movies were acquired of synchronized NRK49F Septin2-eGFP^EN/EN^ cells stained with SiR-tubulin. From these movies, n=206 individual division events were extracted. Only the uninterrupted events (n=129) were used for further analysis. Technical errors included cells leaving the field of view at least partially or floating debris obscuring the analysis. Cytokinetic defects included cells that did not complete abscission or divided into three instead of two daughter cells. Right: Experimental workflow for the acquisition of multiplexed U-ExM images of 17 different targets. U-ExM images were sorted into 6 stages based on the classification of events from the lattice light-sheet movies. By aligning the images along the cytokinetic division axis, average representations were generated: i: averaged mean intensity projections, ii: averaged 3D representations (z-stacks) and iii: averaged axio-radial distribution maps. These representations were used to describe a specific target distribution over pseudotime and to correlate different target distributions with each other. Scale bars are 100 µm in LLS overview images, 10 µm in extracted LLS events, 5 µm in individual U-ExM images, and 1 µm in averaged representations of the U-ExM data.

**Supplementary Figure S5:**
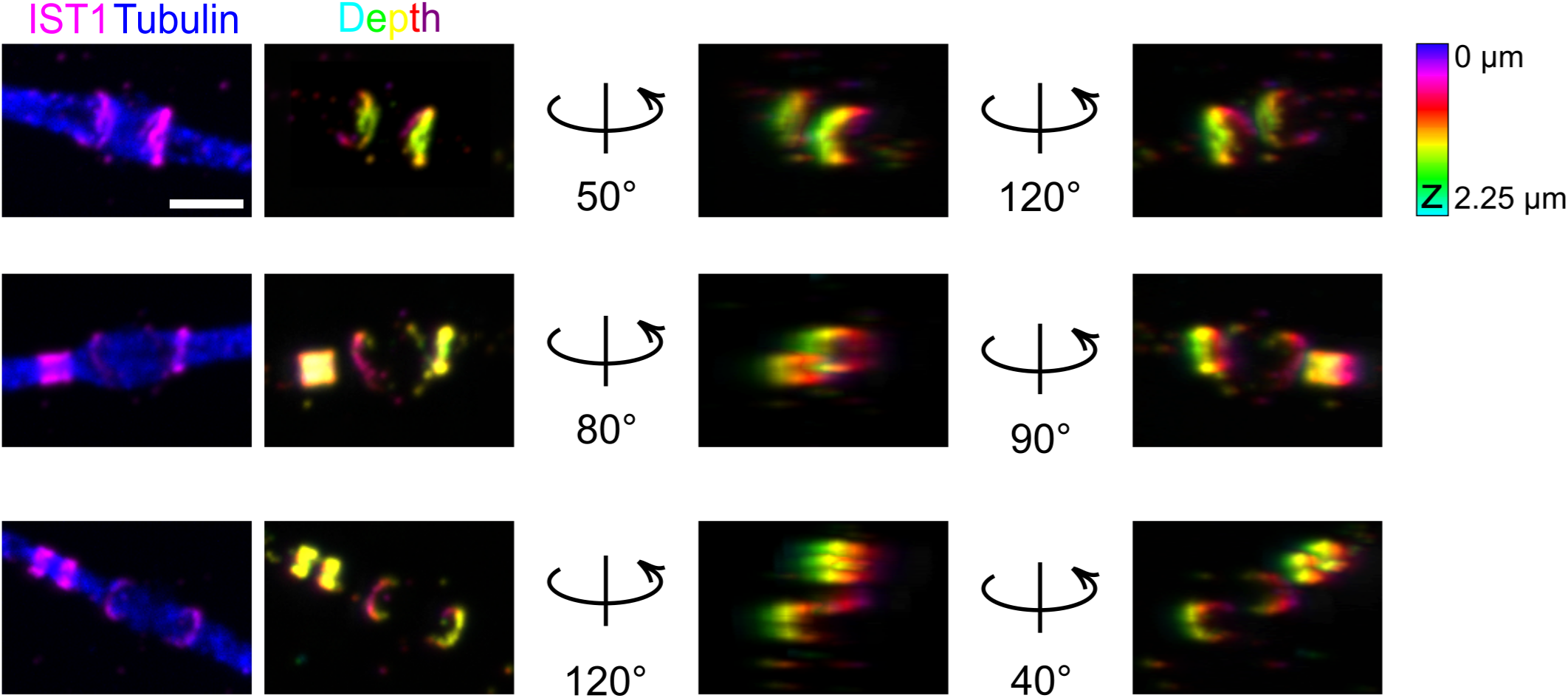
Three dimensional representation of IST1 structures in sub-stages before abscission Three examples of U-ExM volumetric images of IST1 organization. Maximum intensity projections of IST1 (magenta) and tubulin (blue). IST1 signal is color-coded by depth and shown in different rotation angles, emphasizing the ring or cone-like assembly. Scale bar is 1 µm.

**Supplementary Figure S6:**
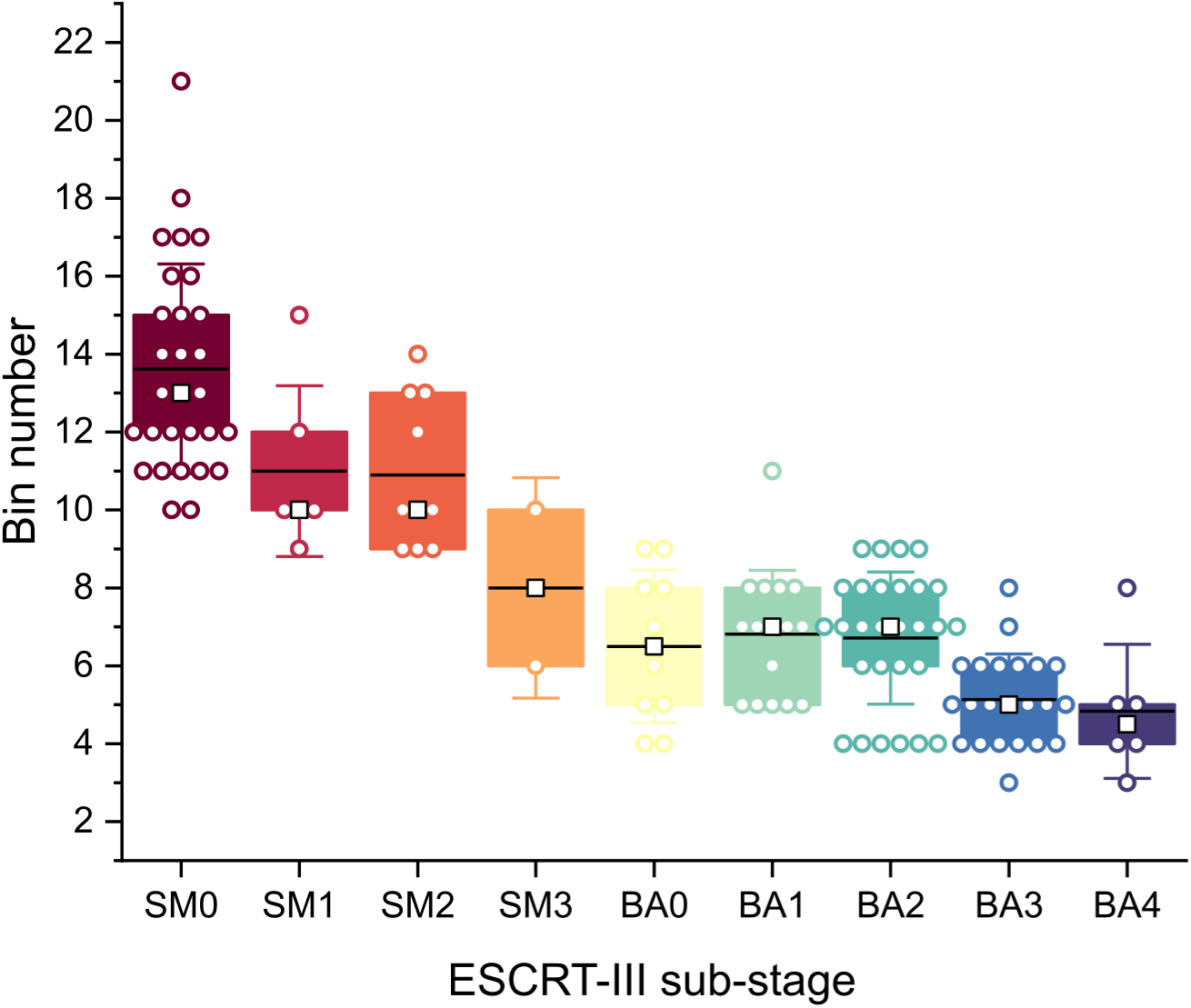
Plot of ESCRT-III assembly and cytokinetic stage vs. pseudotime Sub-stages of ESCRT-III assembly and their PCA-based assignment into pseudotime bins. Lower bin numbers are closer to abscission. Images were assigned based on their tubulin and Septin2-GFP channels. Boxes show 25^th^-75^th^ percentile, black solid line marks the mean, black square marks the median, whiskers show standard deviation.

