## Supplementary Information for "Multiscale imaging of cytokinesis"

List of Supplementary Files

| File | File name | Description |
|---|---|---|
| Supplementary File 1 | pseudotime_images_z-stack_reduced_MTs_sen_DAI_Cek_MK1_Ra1_PR1_CiK_ann_myA_myB_acn_Se7_Se1_Se9_BO4_Tsm_ALm_IS1_CHB_8bit.tif | 5D file including all targets that were analyzed and averaged in this study in the 6 cytokinetic stages (order: RC, CS, RS, SM, BA, A) |
| Supplementary File 2 | pseudotime_images_mean-proj_MTs_sen_DAI_Cek_MK1_Ra1_PR1_CiK_ann_myA_myB_acn_Se7_Se1_Se9_BO4_Tsm_ALm_IS1_CHB_8bit.tif | 4D file, mean projections of all targets that were analyzed and averaged in this study in the 6 cytokinetic stages (order: RC, CS, RS, SM, BA, A) |
| Supplementary File 3 | pseudotime_images_radial-proj_MTs_sen_DAI_Cek_MK1_Ra1_PR1_CiK_ann_myA_myB_acn_Se7_Se1_Se9_BO4_Tsm_ALm_IS1_CHB_8bit.tif | 4D file, axio-radial projections of all targets that were analyzed and averaged in this study in the 6 cytokinetic stages (order: RC, CS, RS, SM, BA, A) |
| Supplementary File 4 | pseudotime_images_mean-proj_MTs_sen_DAI_IS1_CHB_8bit.tif | 4D file, mean projections of the sub-stages of ESCRT-III cone assembly (related to Figure 7) (order: RC, CS, RS, SM0, SM1, SM2, SM3, BA0, BA1, BA2, BA3, BA4, A) |

Abbreviations used for targets in file names (and default order of channels)

MTs tubulin

sen Septin2-GFP

DAI DAPI

Cek CellMask orange

MK1 MKLP1

Ra1 MgcRacGAP1

PR1 PRC1

CiK Citron Kinase

ann Anillin

myA Myosin IIA

myB Myosin IIB

acn actin

Se7 Septin7

| | |
|---|---|
| Se1 | Septin11 |
| Se9 | Septin9 |
| BO4 | BORG4 |
| Tsm | Tsg101 |
| ALm | ALIX |
| IS1 | IST1 |
| CHB | CHMP4B |
